# An *in vitro* quantitative systems pharmacology approach for deconvolving mechanisms of drug-induced, multilineage cytopenias

**DOI:** 10.1101/2019.12.23.886960

**Authors:** Jennifer L. Wilson, Dan Lu, Nick Corr, Aaron Fullerton, James Lu

## Abstract

Myelosuppression is one of the most common and severe adverse events associated with anti-cancer therapies and can be a source of drug attrition. Current mathematical modeling methods for assessing cytopenia risk rely on indirect measurements of drug effects and primarily focus on single lineage responses to drugs. However, anti-cancer therapies have diverse mechanisms with varying degrees of effect across hematopoietic lineages. To improve predictive understanding of drug-induced myelosuppression, we developed a quantitative systems pharmacology (QSP) model of hematopoiesis *in vitro* for quantifying the effects of anti-cancer agents on multiple hematopoietic cell lineages. We calibrated the system parameters of the model to cell kinetics data without treatment and then validated the model by showing that the inferred mechanisms of anti-proliferation and/or cell-killing are consistent with the published mechanisms for three classes of drugs with different mechanisms of action. Using a set of compounds as a sample set, we then analyzed novel compounds to predict their mechanisms and magnitude of myelosuppression. Further, these quantitative mechanisms are valuable for the development of translational *in vivo* models to predict clinical cytopenia effects.

**Author Summary:** Reduced bone marrow activity and levels of mature blood cells is an undesirable side effect of many anti-cancer therapies. Selecting promising lead compounds for further development requires understanding of potential myelosuppressive effects. However, existing preclinical experiments and modeling formulations fail to consider drug effects on multiple blood cell types or the mechanistic differences between how drugs induced myelosuppression. Here we developed a quantitative systems pharmacology (QSP) model that estimates a drug candidate’s effect on multiple precursor and mature blood cell lineages and further distinguishes how the drug affects these populations - through cell-killing or anti-proliferation mechanisms. This modeling formalism is valuable for vetting compounds for therapeutic development and for further translational modeling to anticipate the clinical effects of compounds.

## Introduction

Drug-induced myelosuppression is one of the most severe adverse events (AEs) associated with anti-cancer therapies(1). Myelosupression increases patient fatigue and hinders their daily routines(2, 3), and increases patient risk for infection (4). Understanding patient propensity for AEs is required for clinical optimization of both drug selection and dose schedules. Often anti-cancer therapies specifically optimize efficacy on the basis of minimizing undesirable myelosuppressive effects (5, 6). Despite the frequency of myelosuppression following anti-cancer treatment, predicting the severity of this AE remains challenging(1).

Computational models and pre-clinical experiments remain the standard for anticipating and understanding potential myelosuppressive effects. Predictive toxicology approaches can expedite early phase clinical trials and reduce the number of patients treated with ineffective doses(7). A validation study of 20 compounds demonstrated that pre-clinical *in vitro* measurements of a drug’s 90% inhibition concentrations (IC90) of granulocyte-macrophages was a sufficient predictor of the maximum tolerated dose (MTD) in animals and humans(7). Many modeling approaches have captured the effects of novel compounds on single lineages. For instance, the Friberg model describes the *in vivo* development of neutrophils using multiple transit compartments where drug treatment can affect the self-renewal and proliferation of immature cell types(8). Importantly, these models have supported safety-mitigating strategies in the clinic. Semi-mechanistic modeling combined with clinical data sufficiently captured G-CSF response and neutrophil loss after chemotherapy(9) and identified an optimal blood monitoring schedule during palbociclib treatment(10).

An understanding of mechanistic and lineage-specific effects would advance predictive toxicology approaches. Improved understanding of drug-induced myelosuppression requires a systems-level perspective of hematopoiesis and effects on progenitors to better explain downstream effects on blood cells(11). A challenge to mathematical modeling of myelosuppression is understanding lineage effects in the bone marrow, especially when using indirect measurements in peripheral blood(3,11,12), suggesting *in vitro* measurements will be essential to this advancement. A cell-based assay that analyzed the relative anti-proliferative effects of multiple chemotherapies found that the extent of anti-proliferation was associated with the severity of myelosuppression(13). These findings further suggest that a mechanistic understanding of drug-induced cytopenias can inform vetting of multiple drug candidates.

Modeling effects on multiple lineages and progenitors could be valuable for interpreting differences in toxicity induced by multiple compounds(3, 11), yet advancing predictability requires better mechanistic understanding. For instance, a decrease in neutrophils could be a result of depletion of mature neutrophils or a depletion of granulocyte progenitors. One recent study used rat to human translation to understand how carboplatin-induced DNA damage affected multiple hematopoietic lineages(12). A key feature of their approach was using QSP modeling to learn carboplatin effects on early hematopoietic progenitors in rats and applying this mechanistic understanding to predict clinical rates of cytopenias. They discovered that feedback on multipotent progenitor (MPP) proliferation was insufficient for capturing clinical recoveries, but that adding feedback on MPP maturation could adequately describe clinical data(12). This demonstrates that a mechanistic understanding of cytopenias is valuable for creating meaningful, translational *in vivo* models.

We developed a quantitative systems pharmacology (QSP) model of *in vitro* hematopoiesis (hereafter referred to *in vitro* QSP model) for quantifying the effects of multi-class anti-cancer agents on multiple cell lineages. In contrast to prior modeling work based on *in vivo* studies(12), our model is built upon a set of *in vitro* data generated using a novel multi-lineage toxicity assay (MLTA) and hence has the benefit of reduced animal use and increased throughput. In particular, we first calibrated the system parameters in the QSP model to cell kinetic proliferation data generated in the absence of any drug treatment. We subsequently generated dose-response data for drugs of interest using MLTA and fitted treatment parameters that reflect the extent and dose-dependence of drug effects per lineage. Our motivation was to understand the mechanisms of drug effects, specifically anti-proliferative and cell-killing effects, and the magnitude of these effects on hematopoietic cell lineages, from progenitors to mature cell types. Towards this goal, experimental and computational methods can complement each other, as illustrated in **Figure 1**. While an IC50 value of a drug on a particular cell type can be directly read out from the MLTA treatment data, it represents the cumulative effects on not only the cell type of interest but also all the progenitors that precede it. Through modeling and computational optimization, we can discern the contributing effects on each individual lineage to recapitulate the net observed cell count decrease. Thus, through the deconvolution of the experimental data, the *in vitro* QSP model provides an understanding into mechanistic and lineage-specific drug effects. We tested the model using drugs with known cytopenia mechanisms and used these parameters as sample reference for considering potential cytopenic effects of novel compounds. The method has broad utility for anticipating cytopenic effects and demonstrates the value in using QSP modeling to anticipate potential safety risks in a predictive, and mechanism-driven fashion.

**Figure 1.**
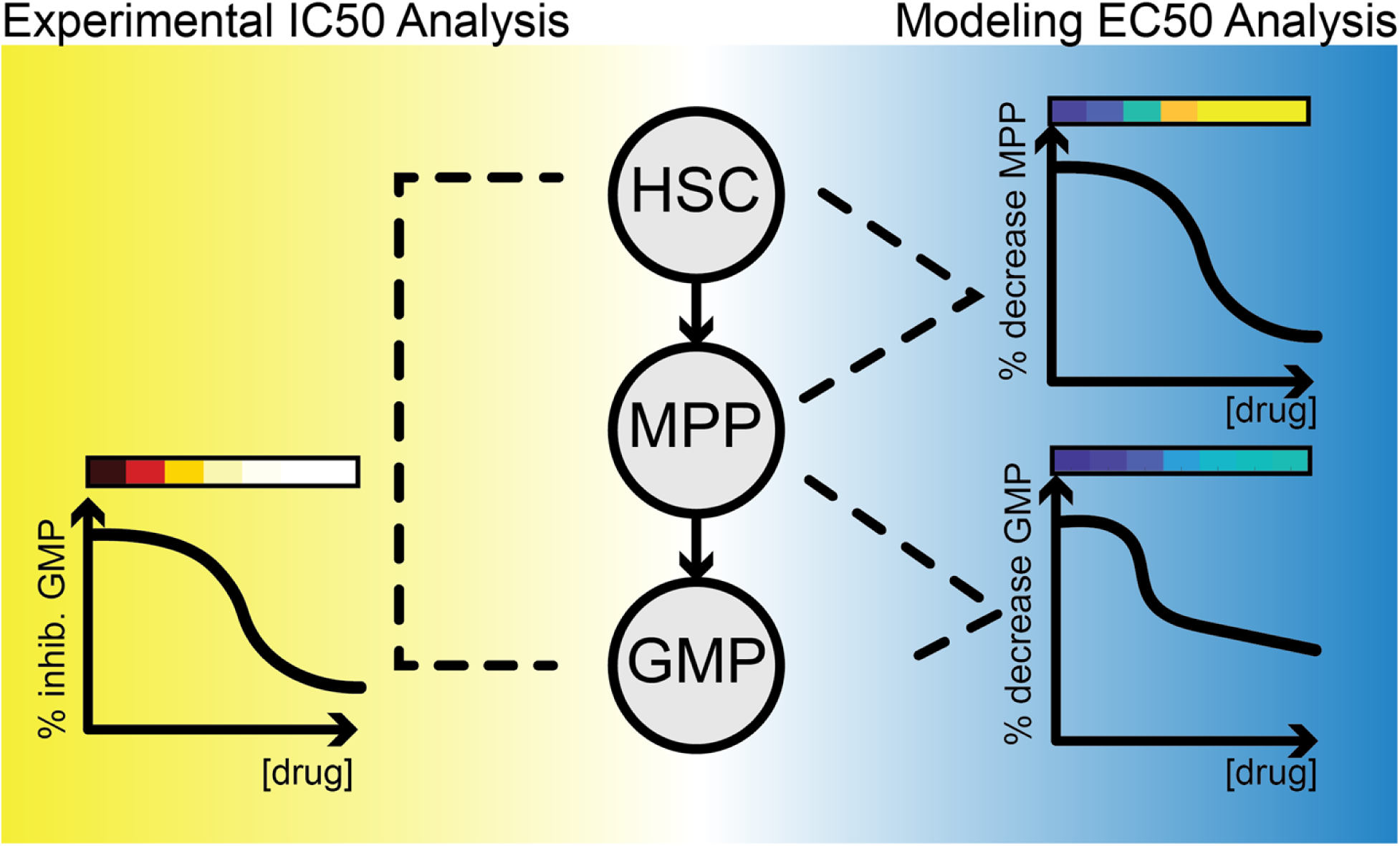
Illustration of the difference between the IC50 value assessed directly from experimental data and model-based deconvolution of mechanisms explaining GMP inhibition. The cartoon shows the percent inhibition of GMP cells at increasing doses of drug. The IC50 value derived from this experimental data represents the cumulative loss along the pathway leading to the formation of GMP, including drug impacts on the early progenitors HSC and MPP. On the right, model-derived Emax and EC50 values represent the drug’s direct effect on cell-killing or anti-proliferation on individual cell types, including HSC, MPP and GMP. In the proposed modeling framework, it is possible to separate effects on a specific cell type, from propagated effects, such as loss of upstream progenitors. The color gradients above all curves represent the percent inhibition (red/yellow color) or percent decrease (blue/yellow color) and these colors are used in later figures.

## Results

### An in vitro QSP model for describing hematopoiesis and cell effects in response to drug treatment

As we endeavored to understand the mechanistic effects of drugs on hematopoietic cell populations, we constructed an *in vitro* QSP model describing hematopoiesis, both in the control condition as well as in the presence of drug treatment (**Figure 2**). The model represents the cell populations measured in the MLTA, their lineage relationships, and the processes of proliferation, differentiation vs renewal, branching and death (see **Figure 2** for the general formulation of the equation system). In **Figure 2A**, the arrows with solid lines denote reactions, whereby the “substrate” leads to the “product” at the end of the arrow; the dashed lines denote the process of proliferation, whereby the same cell type appear as both the “substrate” and “product”. In particular, the model assumes that HSCs replenish themselves, whereas MPPs numbers are maintained by the differentiation of HSCs, as well as the proliferation of MPPs. As **Figure 2A** indicates, the MPPs are assumed to give rise to all of the lineages: erythroid, megakaryocyte, monocyte, granulocyte and lymphocyte branches. Only the most mature cell type of each lineage are assumed to die, shown with red arrows in **Figure 2A**. The model variable “totalViableCells” simply sums up all the live cells represented in the model. The full set of model cell types, parameters, flux equations and ordinary differential equations (ODEs) are further described in (**Supplemental File 1**). Drug effects are modeled at the cell death and proliferation reactions for all cell species (**Figure 2C**). Cell death effects are modeled using an Emax relationship. Anti-proliferation effects are modeled as affecting the basal proliferation rate of the affected cell type.

**Figure 2.**
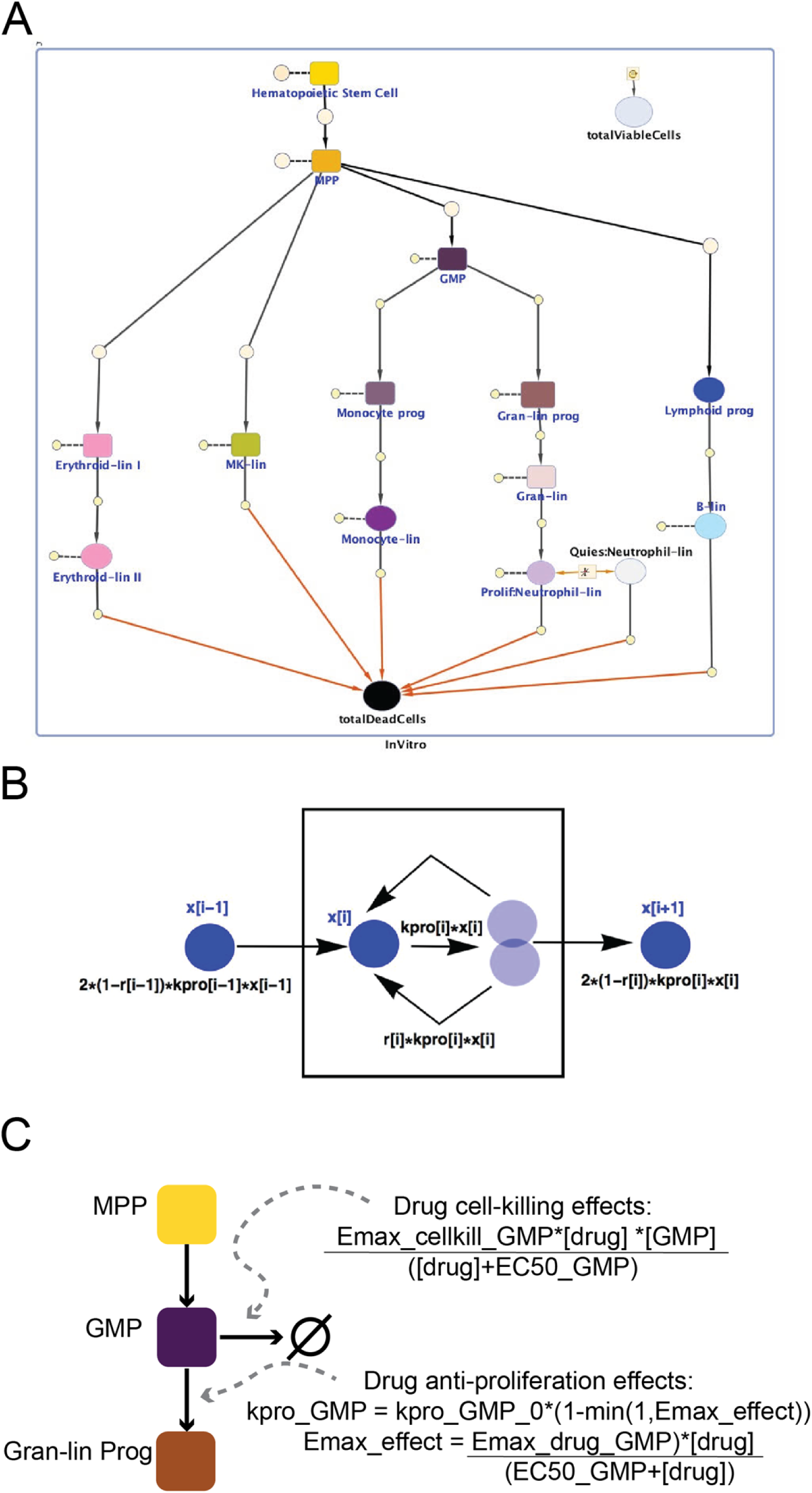
A QSP model for describing in vitro hematopoiesis, cell kinetics, and drug effects. **(A)**The model contains 15 species representing different cell types: hematopoietic stem cells, multi-potent progenitors (MPP), granulocyte-macrophage progenitors (GMP), monocyte progenitors (monocyte prog), monocytes (monocyte-lin), granulocyte progenitors (gran-lin prog), granulocytes (gran-lin), neutrophils, lymphoid progenitors, B-cells (B-lin), megakaryocyte cells (MK lin), early erythroid cells(erythroid I), late erythroid cells(erythroid II), total viable cells (this is the sum of the 13 hematopoietic lineages), and total dead cells. In addition, drug is also represented as a species in the model. The solid arrowed lines in the diagram denote reactions resulting in the formation of products pointed to at the end of the arrow, whereas dashed lines denote reactions whereby the species in connection is both the substrate and product. Under no drug treatment, it is assumed that only the most mature cell types die, as indicated by the red arrows. For the simplicity of illustration, the anti-proliferative and killing effects of drugs on each cell type are not explicitly shown on the diagram, except for the drug-induced killing contributing towards the total number of dead cells. (**B**) The rate of increase in the number of cell type *i*, *x[i]*, depends on the input flux from its predecessor cell type (*x[i-1]*), proliferation flux (*kpro[i]*x[i]*), as well as the fraction renewed (*r[i]*) versus differentiated (*1-r[i]*). Note that drug effects are not represented in this diagram. (**C**) The model captures drug effects at cell death reaction (“Drug cell-killing effects”) and at the proliferation reaction (“Drug anti-proliferation effects”). In this abbreviated schematic we show the drug effects on the GMP cell type and show the example equations for this species.

### The model explains drug-free cell kinetic data

In order to infer the drug effects on various lineages in the multi-lineage toxicity assay (MLTA, publication forthcoming) the proliferation kinetics in the control setting first need to be understood. For this purpose, an experiment was performed to generate time-resolved measurements of cell numbers, at days 0, 2, 3, 4, 5, 6 and 9 (**Figure 3**). The kinetic data was then used to inform parameters in the *in vitro* QSP model. For the purpose of informing model parameters, the data points measured at days 0 and 9 are dropped for model fitting: the reason for dropping data at day 0 is the apparent difference in kinetics between days 0 and 2 (including a decline in counts for certain cell types, potentially due to the cells coming out of liquid nitrogen and hence not recovering until day 2) as compared to the subsequent time points; day 9 data was only generated for exploratory purposes as replating was done after the day 6 measurement, and hence also dropped for the purpose of model fitting. Using the days 2 to 6 data, a hybrid optimization approach (genetic algorithm and local search) was performed to infer a parameter set best compatible with the data (please refer to the Methods section for details). Using the set of parameters identified, the model was able to recapitulate the kinetic data well (**Figure 3**).

**Figure 3.**
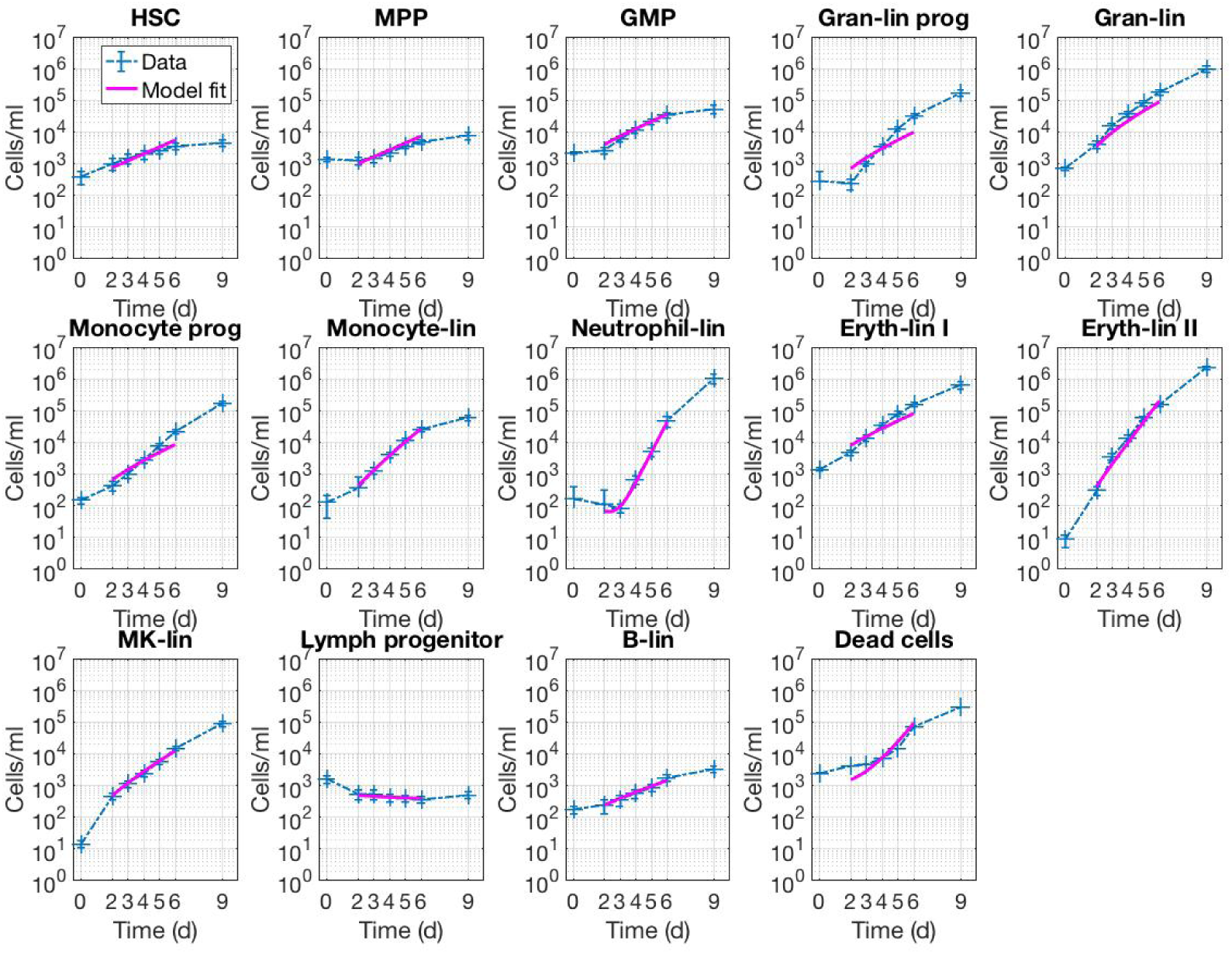
*In vitro* QSP model captures different proliferation kinetics across cell types. Cells/mL are plotted against time (days) for 13 live cell types and the total dead cells. Error bars represent standard error of the mean of six unique donor samples. In all plots, experimental data are shown in blue dashed lines with “+” markers, and model data are shown in solid pink lines.

### Multi-lineage concentration-response data captures cumulative lineage-specific cell responses to drug treatment

We used the MLTA to capture dose-dependent drug effects (publication forthcoming). The 96 well assay used simultaneous differentiation of human donor CD34+ stem cells into multiple lineages and measured these lineages using flow cytometry. We tested 51 compounds (**Supplemental Table 1**). This compound set spanned multiple drug classes and contained drugs with known clinical cytopenic effects as well as novel compounds without clinical data (**Figure 4**).

**Figure 4.**
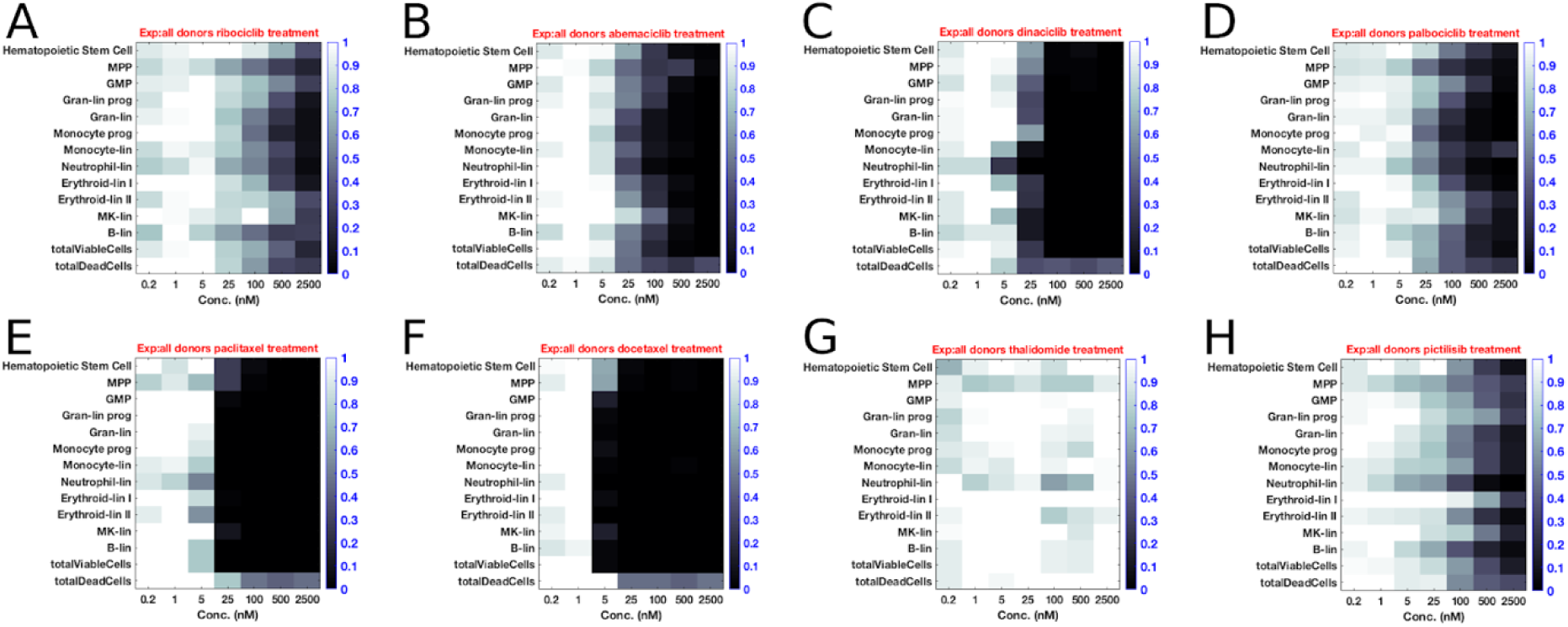
Concentration-response data captures drug effects across multiple lineages. We tested the concentration-response for ribociclib (**A**), abemaciclib (**B**), dinaciclib (**C**), palbociclib (**D**), paclitaxel (**E**), docetaxel (**F**), thalidomide (**G**), and pictilisib (**H**). Extent of shading represents relative cell counts where black and white correspond to fewer or equivalent cell counts relative to vehicle control wells respectively. Doses are in nM concentration and vary across compounds in the full 51 compound set.

The assay captured cumulative changes in numbers across cell types from a sample set of compounds with known hematopoietic toxicities. To provide context for the value of this modeling approach, we have used this sample set throughout the manuscript to describe the analysis and demonstrate usage on a sample of compounds with known hematopoietic effects before applying our method to compounds with unknown effects. Classical chemotherapies exhibited decreased numbers across cell types and was relatively stronger than other drugs in this sample set (**Figure 4, E & F**), and the cyclin-dependent kinase inhibitors (CDKis), ribociclib, abemaciclib, dinaciclib, and palbociclib (**Figure 4, A, B, C, & D**) exhibited relatively more stable cell numbers across lineages. For instance, classic chemotherapies decreased MPP cell numbers to a greater extent and at lower doses than ribociclib or palbociclib. Our negative control, thalidomide (**Figure 4G**), had the least effect on hematopoietic cell types across doses, and the phosphoinositide-3 kinase inhibitor (PI3Ki) pictilisib had a moderate reduction of cells types.

We first applied a traditional, curve-fitting analysis to the concentration response data and identified an IC50 value for each cell type with each drug treatment. We used these IC50 values to plot normalized percent inhibition plots, based on equation 1 (**Figure 5**). Using these relationships, the classic chemotherapies (**Figure 5 E&F**) show a decrease in most cell types and inhibition occurs at lower doses compared to the CDK inhibitors (**Figure 5A-D**). The CDK inhibitors, abemaciclib, dinaciclib, ribociclib, and palbociclib, decreased cell numbers across most cell types, with dinaciclib showing a stronger decrease at lower concentrations. The negative control, thalidomide (**Figure 5G**), did not drastically decrease cell types. The PI3K inhibitor, pictilisib (**Figure 5H**) decreased most cell types. We did PCA analysis on the compounds based on their experimentally-derived IC50 values (**Figure 6**). Compounds were generally well separated by class, and we observed that a single PCA component could explain the variability between compounds. While these plots described observed experimental data well, they did not inform our motivation to understand if the drugs are affecting specific cell types.

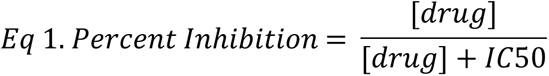

**Figure 5.**
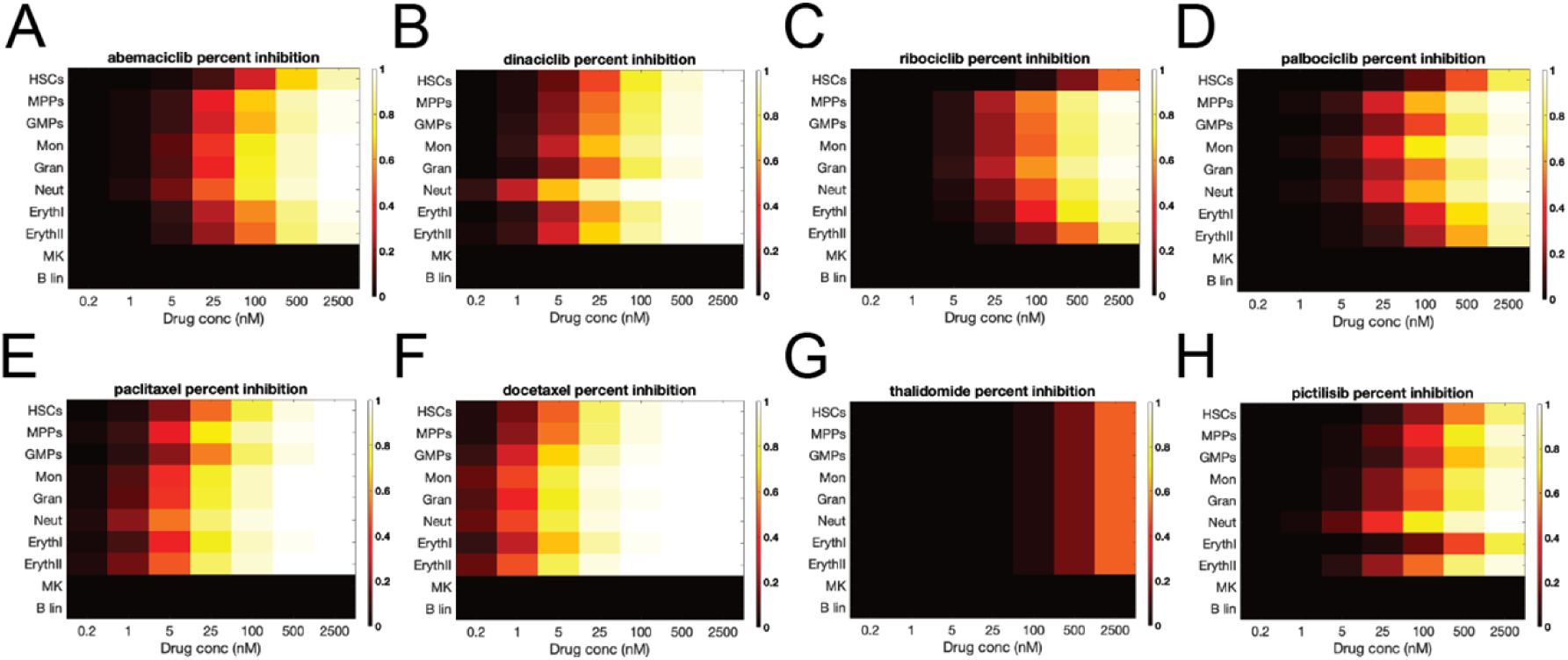
Percent inhibition plotted using IC50 values. Percent inhibition was calculated using IC50 values fitted to the concentration-response data. Black to yellow shading represents increasing inhibition of each cell type: note that as IC50s for MK and B-lin could not be determined from the treatment data, hence they are shown as having 0% suppression. The results indicate that generally a broad set of cell types are inhibited under drug treatment

**Figure 6.**
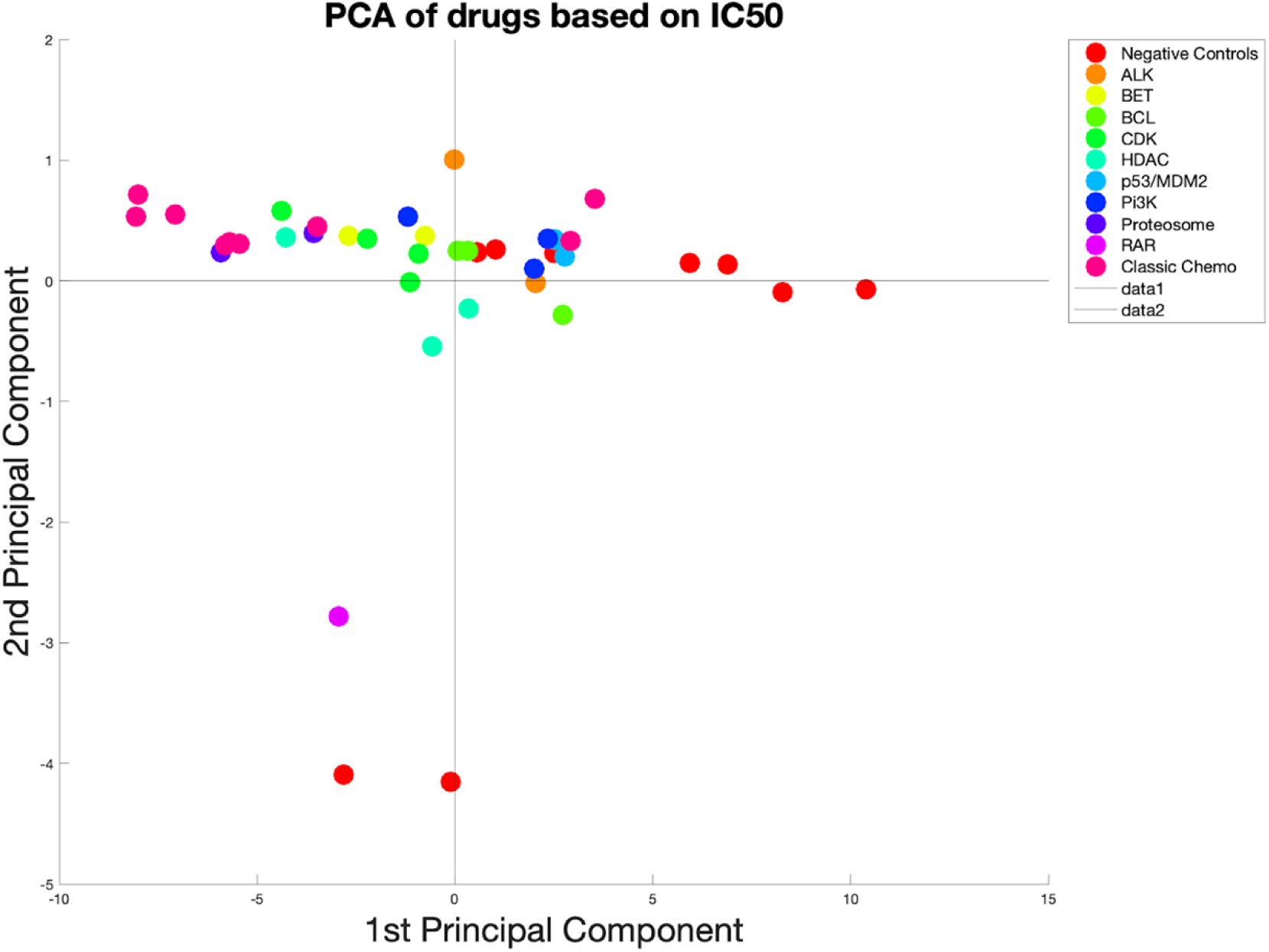
PCA of compounds using experimentally-derived IC50 values. A set of 46 marketed compounds, including 15 negative controls were grouped using principal component analysis. Marker color indicates drug class. The result shows that while almost all the compounds ranged between classic chemotherapies and negative controls on the first principal component, the mechanisms differentiating the compound classes cannot be well identified using the IC50 values alone.

### QSP model fitting identified quantitative, mechanistic hematopoietic effects of drug treatment on each cell type

Using the *in vitro* QSP model, we identified 26 parameters reflecting drug mechanisms for affecting hematopoietic toxicity. We adopted the approach of parsimony in explaining the observed treatment data: namely, for each drug, we fitted a single set of Emax and log(EC50) parameter for each of the 13 cell types measured (Table 1 and full data in **Supplemental Table 2**). Goodness of fit plots and mean squared error across cell types for the sample drug set (highlighted in **Figure 4**) are included in Supplemental Figures 1-8.

**Table 1.**
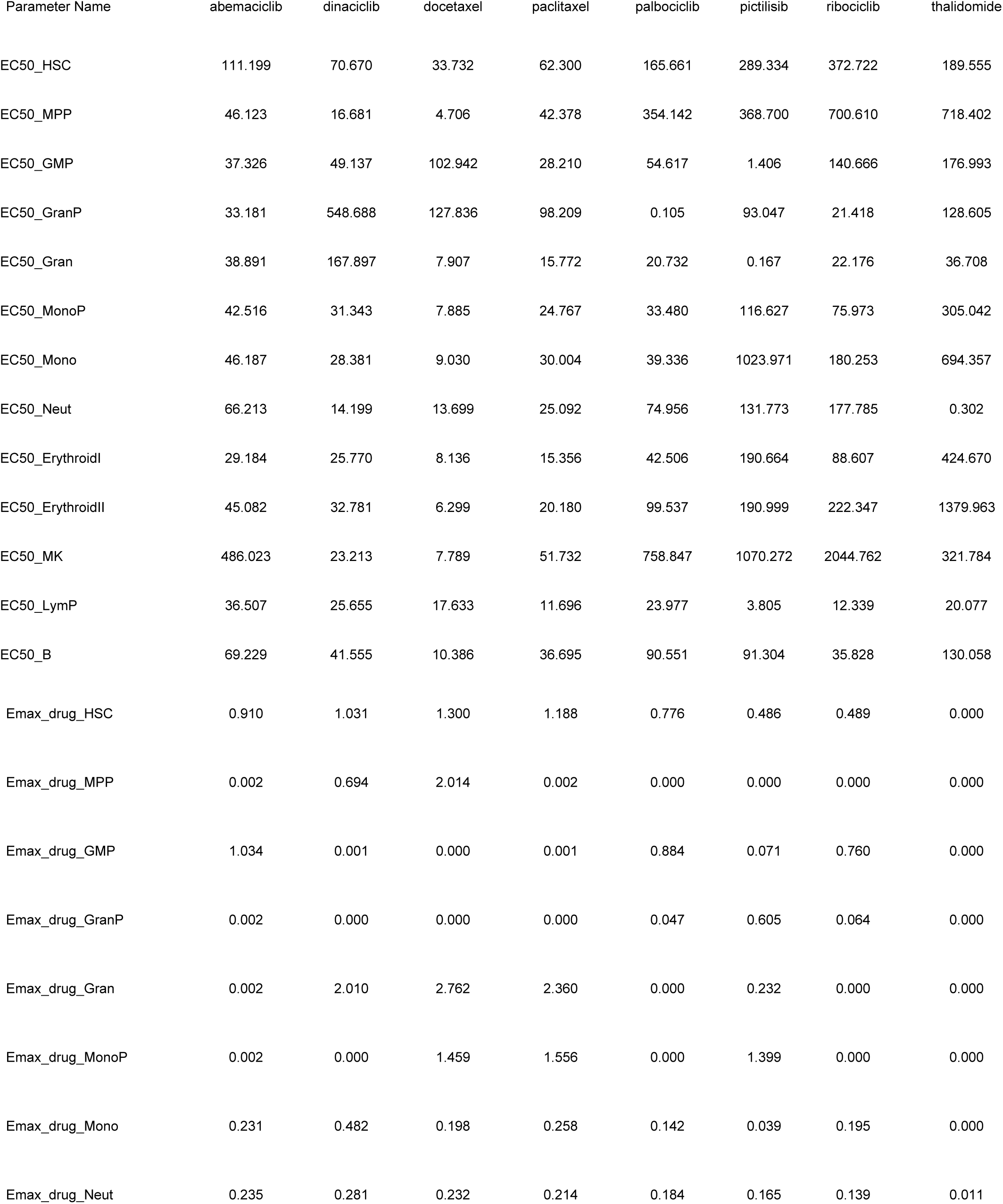
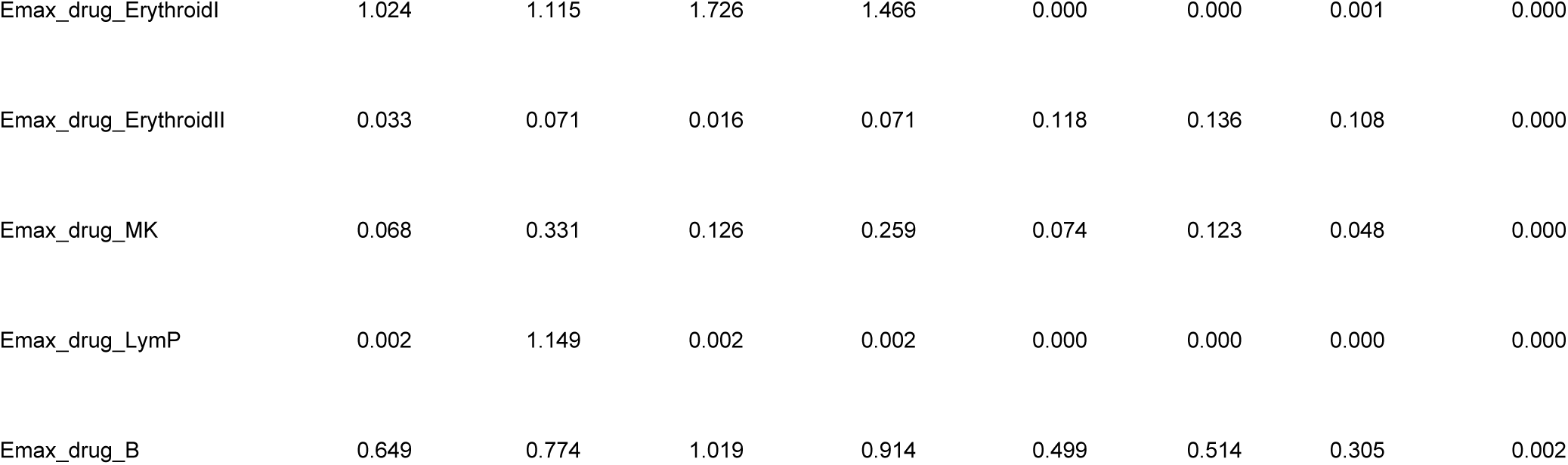
Sample EC50 and Emax parameter values for compounds with known hematopoietic toxicity effects. Thalidomide is included as a negative control and EC50 values above the max tested dose of 2500 nM are extropolated from model fitting.

In this model, we allowed drugs to have both anti-proliferation as well as cell-killing effects. However, in order to describe both effects with the most parsimonious parametric representation, we have developed the following novel formulation. The underlying assumption is that under treatment with increasing concentrations, drugs will first manifest anti-proliferation effects, followed by cell-killing effects at drug concentrations above certain thresholds. The formulation relies upon a single set of Emax and log(EC50): the component of Emax <=1 reflects anti-proliferation of hematopoietic cell lineages and when the drug effect needs to be explained with Emax>1, then the component of Emax larger than 1 reflects cell-killing (Materials and Methods). Conceptually, we interpreted Emax values <= 1.0 to reflect only anti-proliferation effects and Emax values > 1.0 to reflect both complete anti-proliferation combined with cell-killing effects.

As expected, the classic chemotherapies, docetaxel and paclitaxel had relatively strong Emax effects. Specifically, docetaxel induced cell-killing on HSCs (Emax = 1.3), MPPs (Emax = 2.014), granulocytes (Emax = 2.762), monocyte progenitors (Emax = 1.459), and early erythroid cells(Emax = 1.726) and is consistent with published cell-killing effects on hematopoietic progenitors (14). Paclitaxel induced cell-killing on HSCs (Emax = 1.188), granulocytes (Emax = 2.360), monocyte progenitors (Emax = 1.556), and early erythroid cells (Emax = 1.466) and is consistent with the known cytotoxic effects of paclitaxel (15). Comparatively, the CDKis exhibited less cell-killing effects and more anti-proliferative effects. Abemaciclib, dinaciclib, palbociclib, and ribociclib exhibited strong anti-proliferative effects on HSCs (Emax = 0.910, 1.031, 0.776, and 0.489 respectively). Dinaciclib was the only CDKi with a relatively strong cell-killing effect on granulocyte-lineage committed cells (Emax = 2.010). (Table 1 and full data in **Supplemental Table 2**). The estimated anti-proliferative and cell-killing effects of palbociclib and dinaciclib are consistent with published mechanisms(10, 16).

We additionally plotted these results as Emax expressions that illustrate drug effects at increasing concentrations (**Figure 7**). Paclitaxel, docetaxel, and dinaciclib have cell-killing mechanisms (Emax effects > 1.0), though, the classic chemotherapies, paclitaxel, and docetaxel have stronger effects at lower concentrations (**Figure 7**). For comparison, we plotted the equivalent information using the experimental IC50 values (**Figure 5**). As anticipated, the IC50s describe cumulative effects and Emax expressions identify specific cell type effects. In particular, the percent inhibition for palbociclib as shown in **Figure 5D** indicates a drop in cell counts for a block of 8 cell types ranging from HSCs, MPPs through erythrocytes; alternatively, the modeling result shown in **Figure 7D** indicates that palbociclib data can be (parsimoniously) explained by anti-proliferation effects on essentially just the HSCs and GMPs.

**Figure 7.**
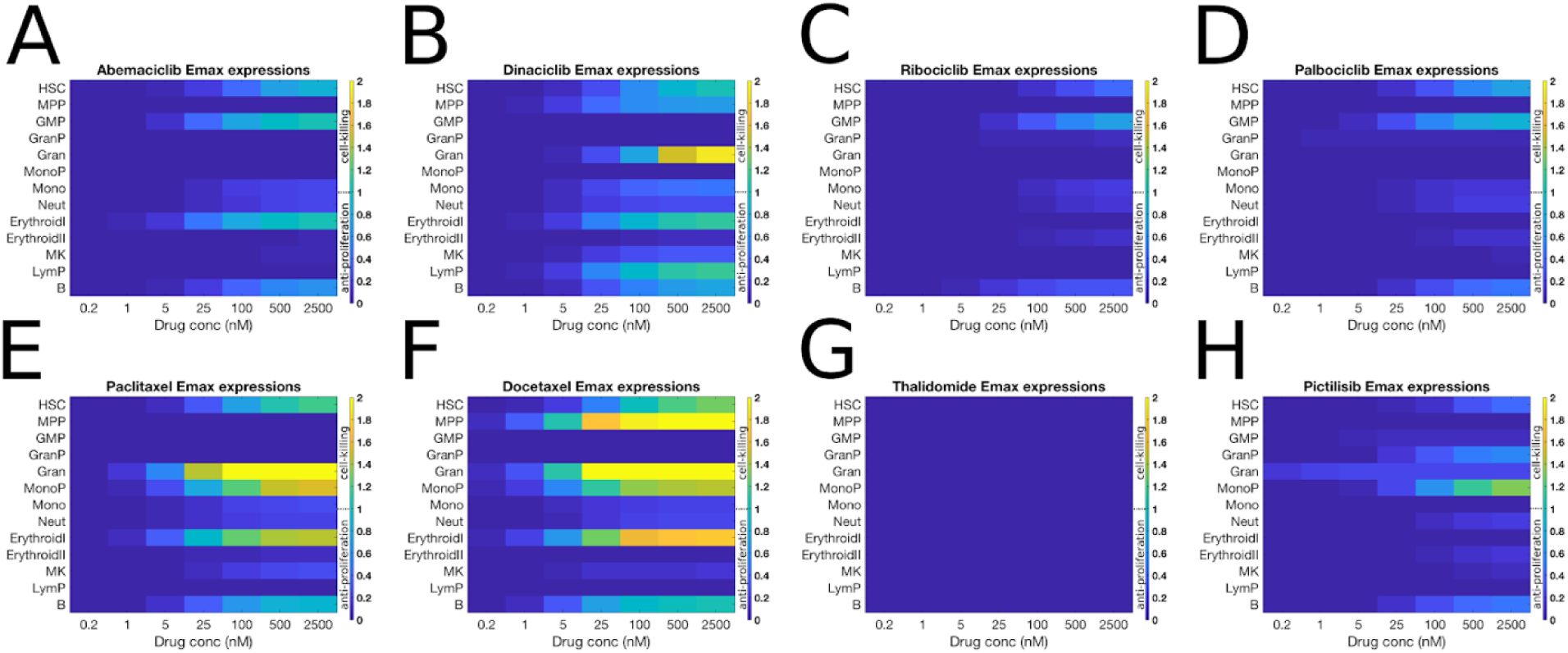
Emax expressions show relative drug effects at increasing concentrations. Emax effects are plotted against concentration for abemaciclib (**A**), dinaciclib (**B**), ribociclib (**C**), palbociclib (**D**), paclitaxel (**E**), docetaxel (**F**), thalidomide (**G**), and pictilisib (**H**). Color bars represent magnitude of Emax effect and concentrations are in nM. The results show that for select targeted therapies, the treatment data can be explained by drug effects on a parsimonious set of cell types.

### Principal component analysis (PCA) separates drugs based on their Emax effects

To consider the potential effects of drugs with unknown clinical toxicity, we used PCA to separate the full 51 drug set based on Emax values (**Figure 9** and Supplemental Figure 9) and EC50 values (Supplemental Figure 10). For this analysis, we used anonymized drug class names to protect molecules in development. When analyzing the EC50 data, we found variance in the data were skewed by compounds with extremely high EC50 values (i.e. compounds that had no detectable EC50 in the concentration range tested). For instance, the parameter log_EC50_Neut had the greatest variance in the data. Many of the class 2 compounds tended to have little to no effect on neutrophils and thus their log_EC_50 values were large and variable. The median and max log_EC50 values for this class were 7.22 (1367 nM) and 8.47 (4754 nM) respectively. Thus, we considered PCA of Emax affects alone as a better formalism for understanding trends among the drugs.

Some classes clustered together, where there was separation between others. For instance, docetaxel and paclitaxel, two microtubule inhibitors from the chemotherapy class, clustered near to each other (**Figure 8**), but separate from the remaining chemotherapies, bortezomib (proteosome inhibitor), cytarabine (DNA synthesis inhibitor), and 5-FU (inhibitor of DNA synthesis). Conversely, dinaciclib was distant from other CDK inhibitors but was proximal to docetaxel and paclitaxel, as compared to all other drugs. Abemaciclib, ribociclib, and palbociclib clustered proximal to each other. The parameters most correlated (> 0.1) with PC1 and PC2 are shown in **Figure 8B**. Emax_drug_Gran, Emax_drug_ErythroidI, and Emax_drug_GMP were relatively well-correlated to principal components 1 and 2 as compared to the Emax values for other cell types (**Figure 8**) and had the greatest variance across the compounds. The correlation of these parameters likely explains the proximity of paclitaxel, docetaxel, and dinaciclib in PCA space because these compounds shared similar Emax parameters for granulocytes, early erythroids, and GMPs. The coefficients for the remaining unlabeled variable names are included in **Supplemental Table 3**.

**Figure 8.**
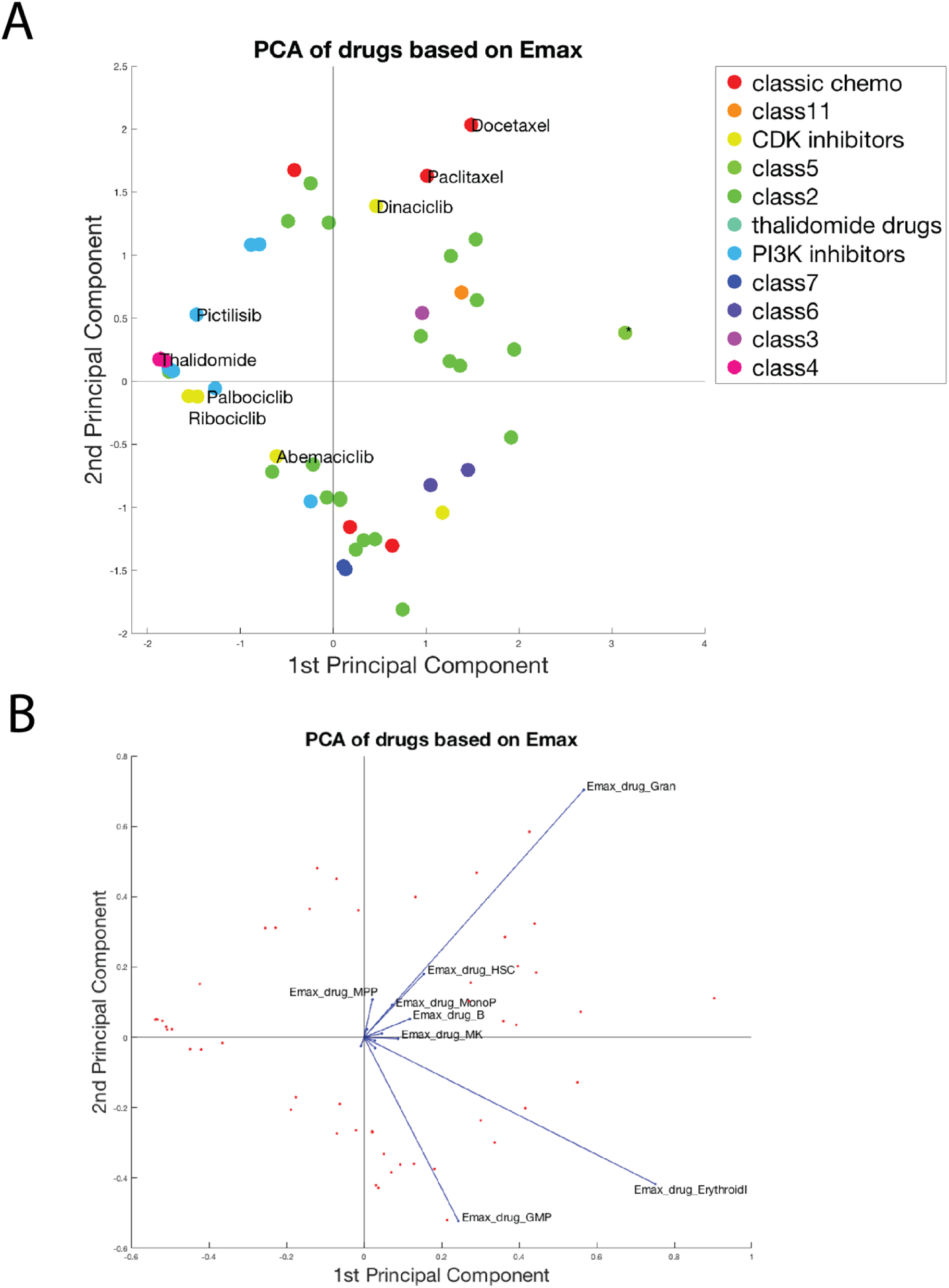
PCA separates drugs based on Emax effects. The 51 compounds are plotted in PCA space (**A**). Marker color corresponds to drug class. Drugs and variables contributing to the top two components are plotted in PCA space (**B**). Note: in figure **A** there is only one drug in class 5 and it is marked with an * to distinguish this compound from the remaining class 2 drugs.

**Figure 9.**
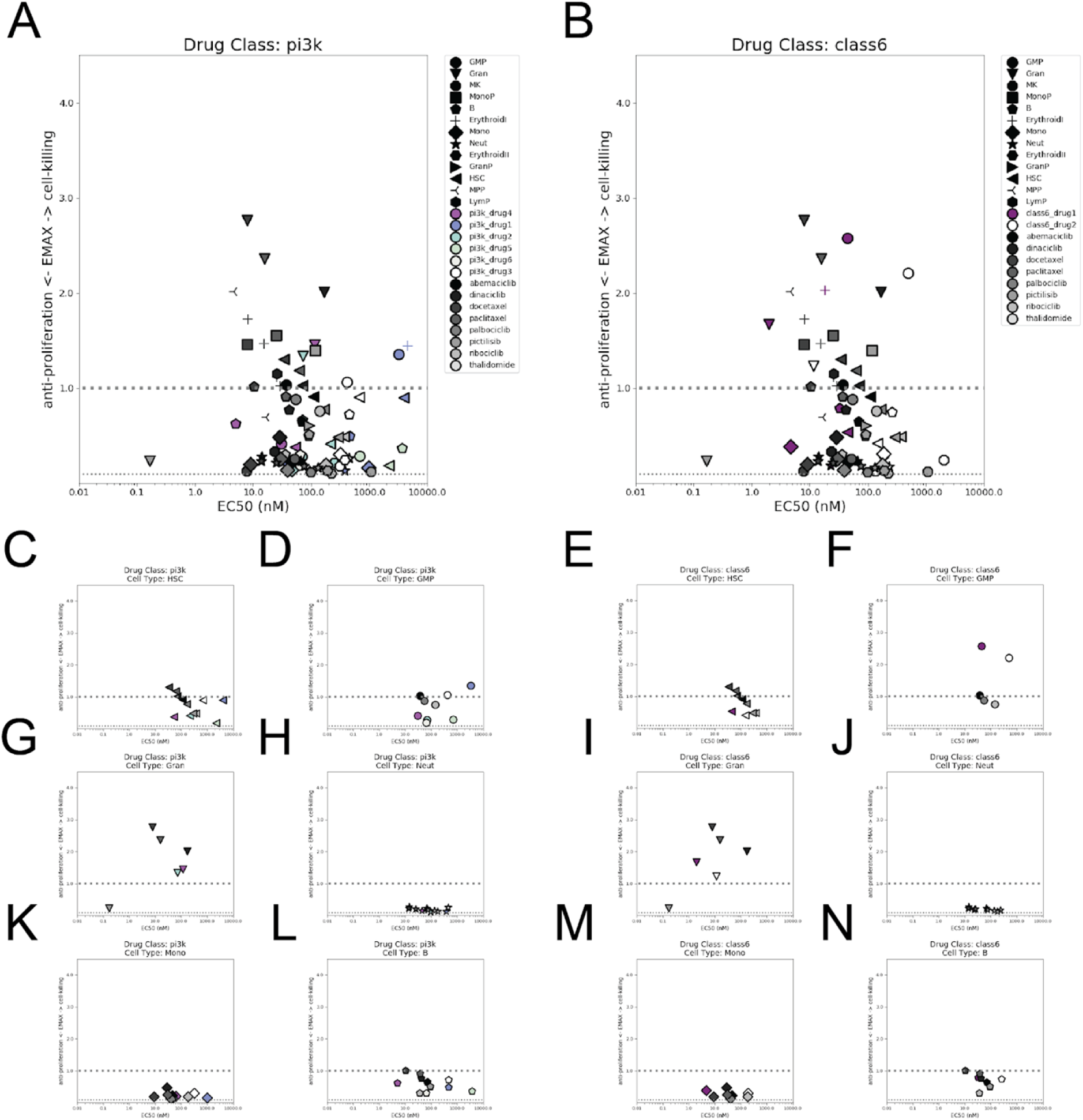
Mechanistic effects of developmental compounds per cell type compared to the sample set. Emax parameters per cell type are plotted against the EC50 values for each cell type. Marker shape represents cell type and marker shading represents either the sample set (black gradient, all figures), PI3K inhibitors (multi-color, **A**), or class 6 (purple, **B**). The same data are broken into cell type plots for PI3K inhibitors: HSCs, GMP, granulocytes, neutrophils, monocytes, B-cells (**C,D,G,H,K,L**); or for class 6 drugs: HSCs, GMP, granulocytes, neutrophils, monocytes, B-cells (**E,F,I,J,M,N**). In all plots, the dashed line represents where Emax = 1.0. Only a subset of cell type plots are shown. Note: only compounds with Emax effects > 0.1 are plotted; compounds such as the negative control, thalidomide, had no effects above 0.1 and are not shown.

### Analysis of mechanistic effects enables consideration of experimental drug effects

To further scrutinize novel compounds for potential hematological toxicity, we considered their magnitude of effect per cell type relative to the sample set of compounds considered in **Figure 4**. (**Figure 9**, and supplemental Figure 11). Drugs in class 3 and class 6 (**Supplemental Figure 11B** and **Figure 10B**) had relatively similar Emax effects compared to the sample set and these effects were shifted to higher EC50 values. The single drug in class 5 had strong cell-killing effects on early erythroid cells and GMP cells, similar to dinaciclib, docetaxel, and paclitaxel (**Supplemental Figure 11D**, triangles and diamonds respectively). The class 4 compounds (**Supplemental Figure 11C**) had relatively little to no Emax effects compared to the sample set. The PI3K inhibitors had similar anti-proliferative but reduced cell-killing effects as compared to the sample set (**Figure 9A**). The remaining named compounds tested (**Supplemental Figure 11E**) had comparable Emax effects relative to the sample set.

## Discussion

Here we presented an approach using a QSP model of *in vitro* hematopoiesis for learning lineage-specific drug mechanisms of myelosuppression. Our ordinary differential equations model described cell kinetics in the absence of drug treatment, as well as drug effect parameter values Emax and EC50 that explain the observed drug effects on multiple hematopoietic cell types. The major innovation in our approach is the deconvolution and mechanistic interpretation of drug effects on multiple hematopoietic lineages including early progenitors and mature blood cells. By this modeling approach, we were able to estimate the drug effect parameters (Emax and EC50) on each cell type in the hematopoiesis pathway in a manner that is independent of study treatment duration and enables model-based *in vitro* to *in vivo* translation that is underway. We expect the model-based translational approach to be more predictive of clinical outcomes, due to the fact that known differences between *in vitro* and *in vivo* hematopoiesis are captured in the mathematical models (including proliferation kinetics and feedback mechanisms mediated by cytokines such as G-CSF, EPO and TPO).

We used these parameters with multivariate and comparative analyses across compounds to anticipate if and the extent of hematological toxicity of novel compounds. More specifically, we selected an example set and compared parameters of novel compounds in development to the parameters associated with the sample set. We anticipated this modeling and interpretation paradigm to be valuable for early stage development decisions. Multivariate analysis enabled comparison of novel compounds based on their complete set of Emax values. Here we highlighted that dinaciclib is broadly more similar to the classic chemotherapies than to other CDK inhibitors. Further, cell-type-specific analysis of drug classes compared to the sample set enabled cell-type specific consideration of within-class drugs. This analysis could provide support for discriminating one drug class over another, or rank-order compounds within the same class based on their toxicity profiles. For instance, the PI3K and class 4 compounds had favorable toxicity parameters compared to the sample set where the single agent in class 5 had much stronger cell-killing effects than the sample set. The platform is flexible, and the sample drug set can be modified to reflect compounds and toxicity effects specific to a development program.

Many compounds are known to be cytotoxic to hematopoietic cells, yet, cytopenias measured in the blood do not reflect the full extent of damage to the hematopoietic system(17). Understanding of mechanisms for myelosuppression would further inform first-in-human trials and help anticipate possible adverse-event mitigating strategies such as appropriate cytokine therapy by increasing evidence-based practices (3, 11). For instance, cytokine therapy is already employed for patients receiving anti-cancer therapy. Understanding how and where the drug affects the hematopoietic lineages will help inform whether or not a specific cytokine therapy might mitigate potential safety concerns; for instance, a drug that results in the loss of hematopoietic stem cells would have a different risk implication from one that depletes the granulocyte lineage alone.

A mechanistic understanding of lineage-specific drug effects could inform further translational modeling. Specifically, we are actively developing an *in vivo* clinical model that translates *in vitro* drug parameters to simulate multiple cytopenias, including thrombocytopenia, neutropenia, and anemia. This *in vivo* model could inform optimal dosing schedules considering multiple cytopenia effects and ultimately inform personalized treatment approaches (18).

## Materials, Methods, and Model

### In vitro multilineage hematopoietic toxicity assay culture system

A custom optimized cytokine cocktail was developed using commercially available cytokines (Peprotech, New Jersey) to facilitate simultaneous multi-lineage differentiation and self-renewal of freeze-thawed primary human bone marrow-derived CD34+ cells in SFEM II media (cells and media from StemCell Technologies, Vancouver). Cultures were carried out in ultra-low attachment 96-well plates (Corning, New York) with test article added on day 0 and cultured at 37°C, 85% relative humidity, and 5% CO2 for six days before interrogating drug impact via 17-parameter flow cytometry.

### Flow cytometry panel and analysis

The flow cytometry panel used to define hematopoietic subsets was composed of CD34-PECy7, CD90-PECY5, CD38-AF488, CD371-PE, CD45Ra-BV421, CD14-BV605, CD42b-AF647, CD10-PEDazzle594, CD235a-PerCPCy5.5 (BioLegend, California), CD15-BUV395, CD41a-APCH7, CD71-BV786 (BD Biosciences, California), and DAPI for dead cell exclusion (ThermoFisher, California). In short, Day 6 cells were harvested, FcR-blocked (FcR binding inhibitor, ThermoFisher), surface-stained, then analyzed in a buffer containing TruCount Control Beads using a BD LSR Fortessa SORP fitted with a high-throughput sampler (BD Biosciences, California). Data was analyzed in Diva 6.0.2 and exported to CSV where after all cell data was transformed to cell per mL then normalized on a per-donor basis to untreated control wells to determine drug impact.

### Measuring drug-free cell kinetics

Six replicate samples of human donor CD34+ stem cells were seeded in 96 well plates and exposed to a customized cytokine cocktail to promote differentiation (publication forthcoming). Cells were maintained in culture for six days and cell lineages were measured using flow cytometry as described above.

### Model simulation and calibration to cell kinetic data without treatment

We implemented the model using the Simbiology toolbox in Matlab, version 2019A. The flux equations, ODEs, model parameters and model cell types are further described in **Supplemental File 1**. The model tracks 13 live cell types: hematopoietic stem cells (HSCs), multi-potent progenitors (MPPs), granulocyte-macrophage progenitor (GMPs), granulocyte progenitor (Gran-lin prog), granulocytes (Gran-lin), monocyte progenitors (monocyte prog), monocytes (monocyte-lin), neutrophils (neutrophil-lin), early erythroid cells(eryth I), late erythroid cells (eryth II), MK cells (MK-lin), lymphocyte progenitors (LymP) and B cells (B-lin); the total number of dead cells is also described (totalDeadCells). The ODE system describes the various steps of differentiation, proliferation, and renewal of hematopoietic stem cells into mature lineages. The most mature cell type of each lineage (that is: eryth II, MK-lin, monocyte-lin, neutrophil-lin and B-lin) are assumed to die, at a uniform rate (kDeath). In summary, the system parameters of the model include cell-type specific renewal rates (r[i]), proliferation rates (kpro[i]), branching rates (kbranch[i]) and a uniform rate (kDeath).

The 27 system parameters of the model as well as the initial cell counts (at time=0) for each of the 14 cell type modeled were fitted to describe the mean of the kinetic data over the 6 donors. In particular, a hybrid optimization approach combining genetic algorithm with local search was used (as implemented in the Matlab function, ga) to obtain an optimal parameter set that best explain the experimental data. For the algorithm settings, the maximum number of generations (MaxGenerations) and the population size (PopulationSize) were set to 50 and 500 respectively. Appropriate lower and upper bounds ([LB, UB]) were set on the parameter values and initial conditions, as follows:

- Renewal rates: [0.5, 1] for HSCs and [0, 0.5] for all other proliferating cell types
- Branching rates: [0.001, 1] and summing up to 1 at each branching step
- Proliferation rates: [0, 4/log(2)]
- Death rate: [0, 2]
- Initial conditions: [0.1, 1] x measured cell counts @ t=0

### Multi-lineage toxicity assay (MLTA) using anti-cancer therapies

Donor material was collected and cultured as described above. The number of replicate donor samples per drug ranged from 2-7 and varied per drug **Supplemental Table 1**. Drug treatment was added a time 0 and cells were maintained in culture for 6 days. Concentrations tested for each drug compound are contained in **Supplemental Table 1**. Following treatment, cell populations were measured using flow cytometry. We extracted cell counts from flow cytometry data, normalized to bead counts, and corrected for well volume.

### Data normalization and pooling across donors

Each well was normalized to the average of six vehicle control wells. For drugs that were tested across multiple runs of the assay, vehicle-normalized donor data were pooled, and we used the average of these pooled data for further modeling.

### Creating an in vitro model for hematopoiesis in the presence of drug treatment

We implemented drug effects using an Emax model for the drug’s effect on each cell lineage. The 13 Emax parameters (Emax_drug_#) are unitless and can vary between 0 and 2 and where # represents one of the previously described 13 cell types. We interpreted Emax values less than or equal to one as anti-proliferative, and Emax values greater than one as indicating a cell-killing mechanism. The relationships for these effects are included in the model using the following equations:

#### Anti-proliferation

kpro_# = kpro_#_0*(1-min(1,Emax_drug_#)*drug/(exp(log_EC50_#)*EC50_unit+drug))

#### Cell-kill

(Emax_cellkill_#*drug)/(drug+exp(log_EC50_#)*EC50_unit))*[#], whereEmax_cellkill_# = max(0,Emax_drug_#-1)*one_over_day

In the above equations, *one_over_day* and *EC50_unit* have units of (1/day) and (1 nM) respectively to maintain correct dimensionality and [#] represents the concentration of the relevant cell type. The first equation describes the drug antiproliferative effect in the net proliferation rate for a given cell type. The second equation describes the cell-killing Emax effect.

When developing the model, we considered different formulations: (a) weighting the objective value to prioritize dead cells (b) using a regularization parameter to encourage a parsimonious solution. We assessed the contributions of these formulations using mean-squared error and goodness of fit plots for drugs with known mechanisms. We discovered that weighting the distance between the observed and estimated value for the dead cell populations by a factor of two was necessary to fit known cell-killing and anti-proliferation mechanisms. Further, we used a regularization parameter of 0.1 to penalize parameters that were close to zero and ultimately produce a parsimonious set of parameters to explain drug effects.

### Extracting EC50 and EMAX effects to explain drug myelosuppression mechanisms

For estimating mechanistic parameters, we used optimization to fit normalized concentration-response data for each drug across 13 cell types: hematopoietic stem cells (HSCs), multi-potent progenitors (MPPs), granulocyte-macrophage progenitor (GMPs), granulocyte progenitor (Gran-lin prog), granulocytes (Gran-lin), monocyte progenitors (monocyte prog), monocytes (monocyte-lin), neutrophils (neutrophil-lin), early erythroid cells(eryth I), late erythroid cells(eryth II), MK cells (MK-lin), and B cells (B-lin), Optimization identified a set of 26 parameters: 13 total Emax parameters and 13 total EC50 parameters (one Emax and one EC50 parameter per cell lineage).

### Principal component analysis (PCA) of 51 compound set

We conducted PCA of the drug parameters estimated from the 51 compounds using Matlab version 2019A (selecting the singular value decomposition algorithm and other default settings), with the aim of reducing the dimensionality of the inferred parameters and help identify patterns within compound classes. Because all of our Emax values and log_EC50 values were on the same scale, our analysis required no further data scaling. We tested both PCA using only Emax effects and only log_EC50 parameters.

### Mechanistic plots of lineage specific drug effects

We created cell-type specific plots using Python version 2.7.16.

## Acknowledgements

The authors would like to thank Brendan Bender, Dale Miles, Jin Jin, Saroja Ramanujan, and Amita Joshi for their discussions and helping to make this work possible.

## Supplementary File Legends

*supplementary_table1_drugs_and_doses.xlsx:* This contains drug names or anonymized names and the doses tested in the MLTA. This is the raw data for the paper.

*supplemental_table2_drug_emax_logEC50_params.xlsx:* For all drugs, this contains parameters fitted from the QSP model.

*supplemental_table_3_coefficients_from_PCA.xlsx:* This contains coefficients from the PCA analysis for all drugs.

## Supplemental Material

**Supplemental Figure 1.**
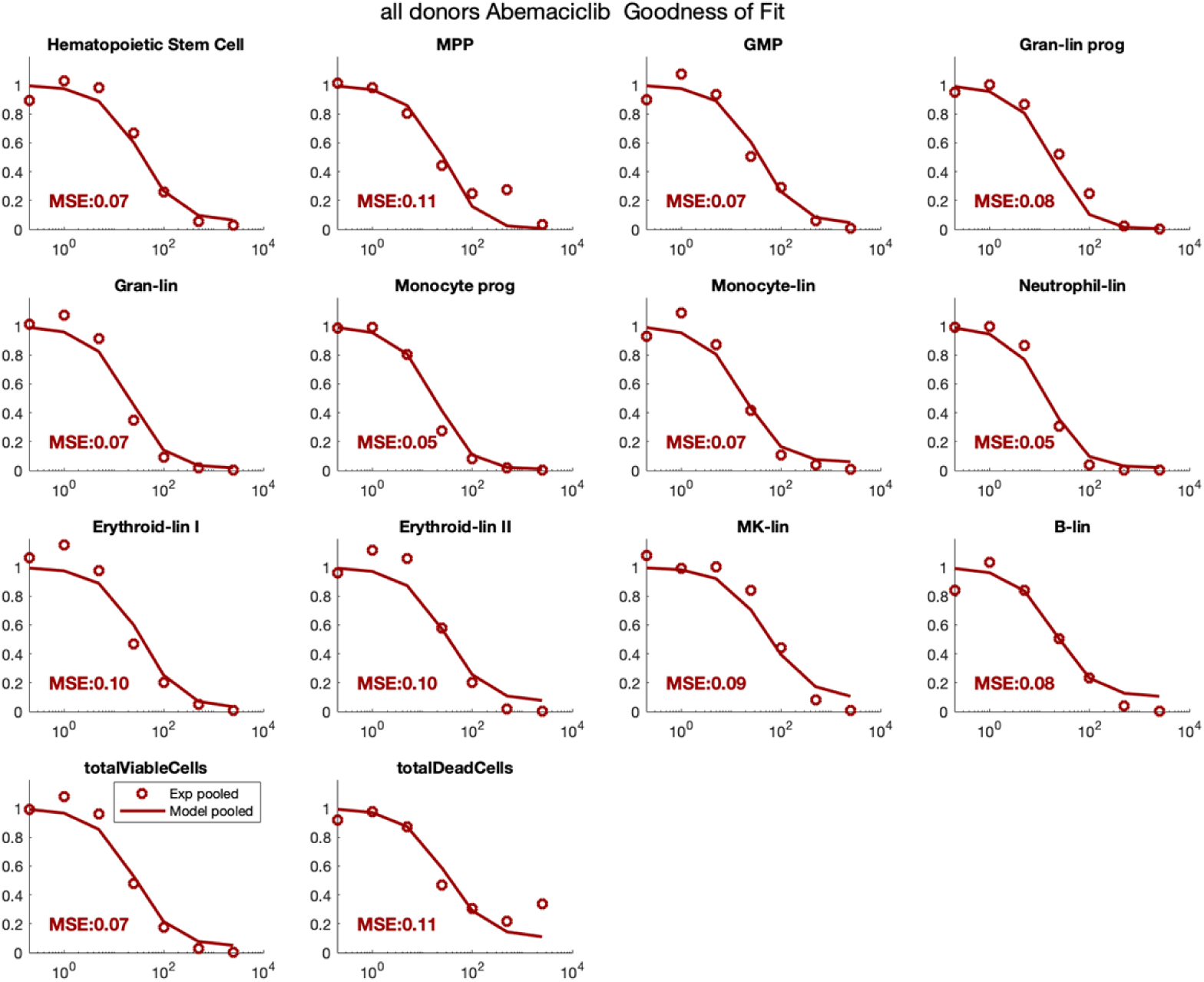
Goodness of fit across cell types for abemaciclib. For each cell type and total live and viable cells, normalized cell counts from experimental data (open circles) and simulated results (solid line) are plotted against concentration (nM). Additionally, each plot includes the mean squared error of the difference between experimental and simulated data.

**Supplemental Figure 2.**
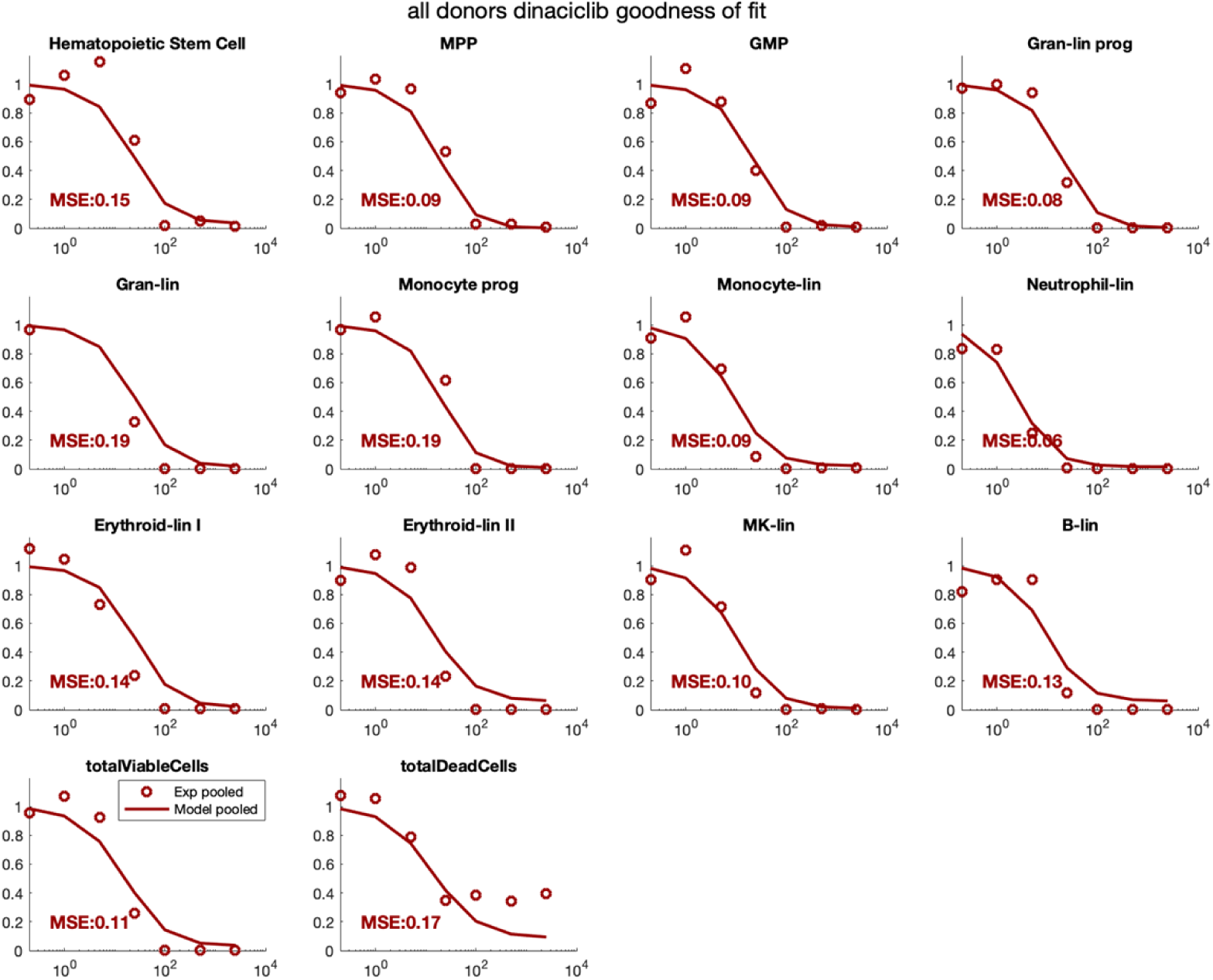
Goodness of fit across cell types for dinaciclib. For each cell type and total live and viable cells, normalized cell counts from experimental data (open circles) and simulated results (solid line) are plotted against concentration (nM). Additionally, each plot includes the mean squared error of the difference between experimental and simulated data.

**Supplemental Figure 3.**
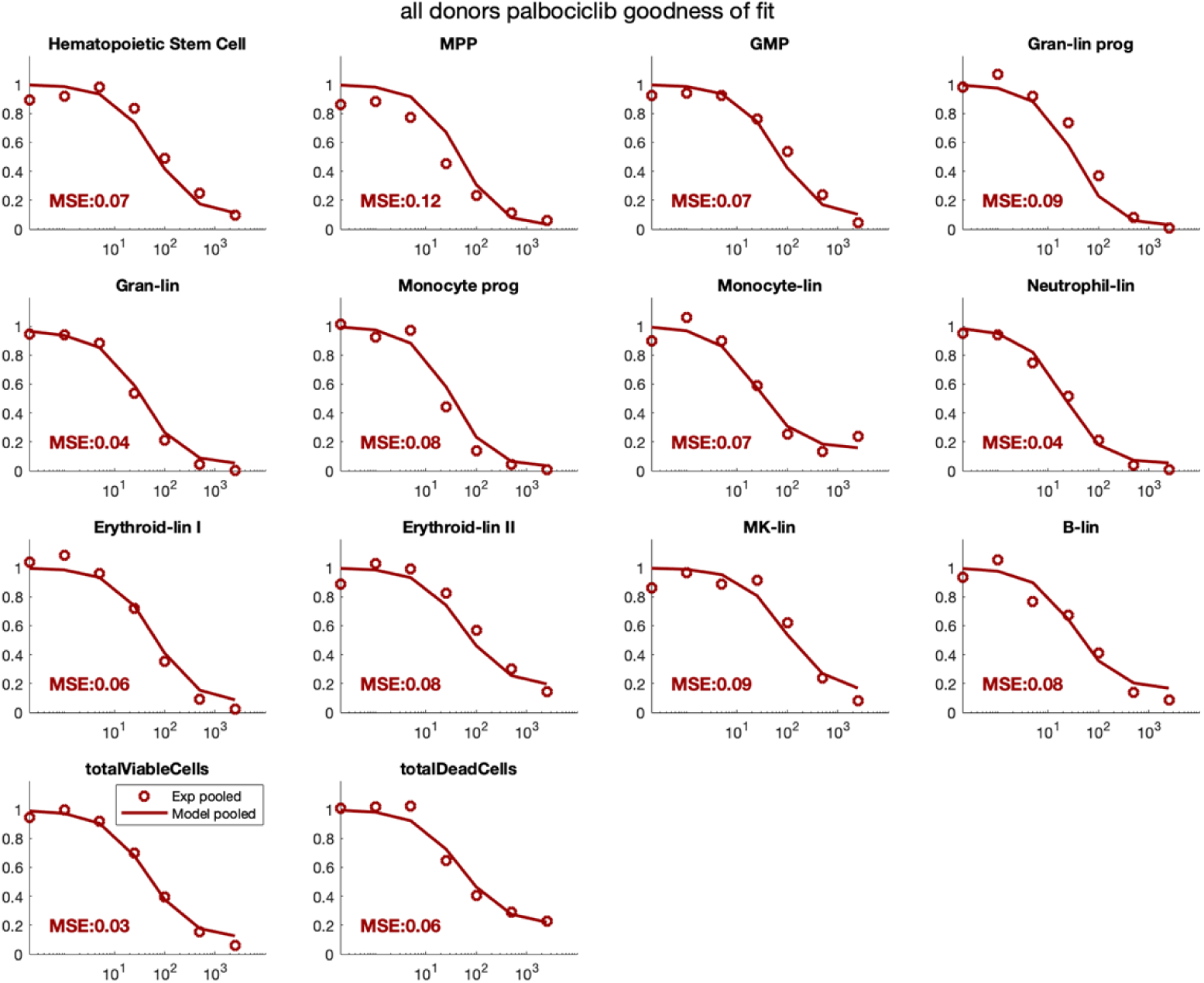
Goodness of fit across cell types for palbociclib. For each cell type and total live and viable cells, normalized cell counts from experimental data (open circles) and simulated results (solid line) are plotted against concentration (nM). Additionally, each plot includes the mean squared error of the difference between experimental and simulated data.

**Supplemental Figure 4.**
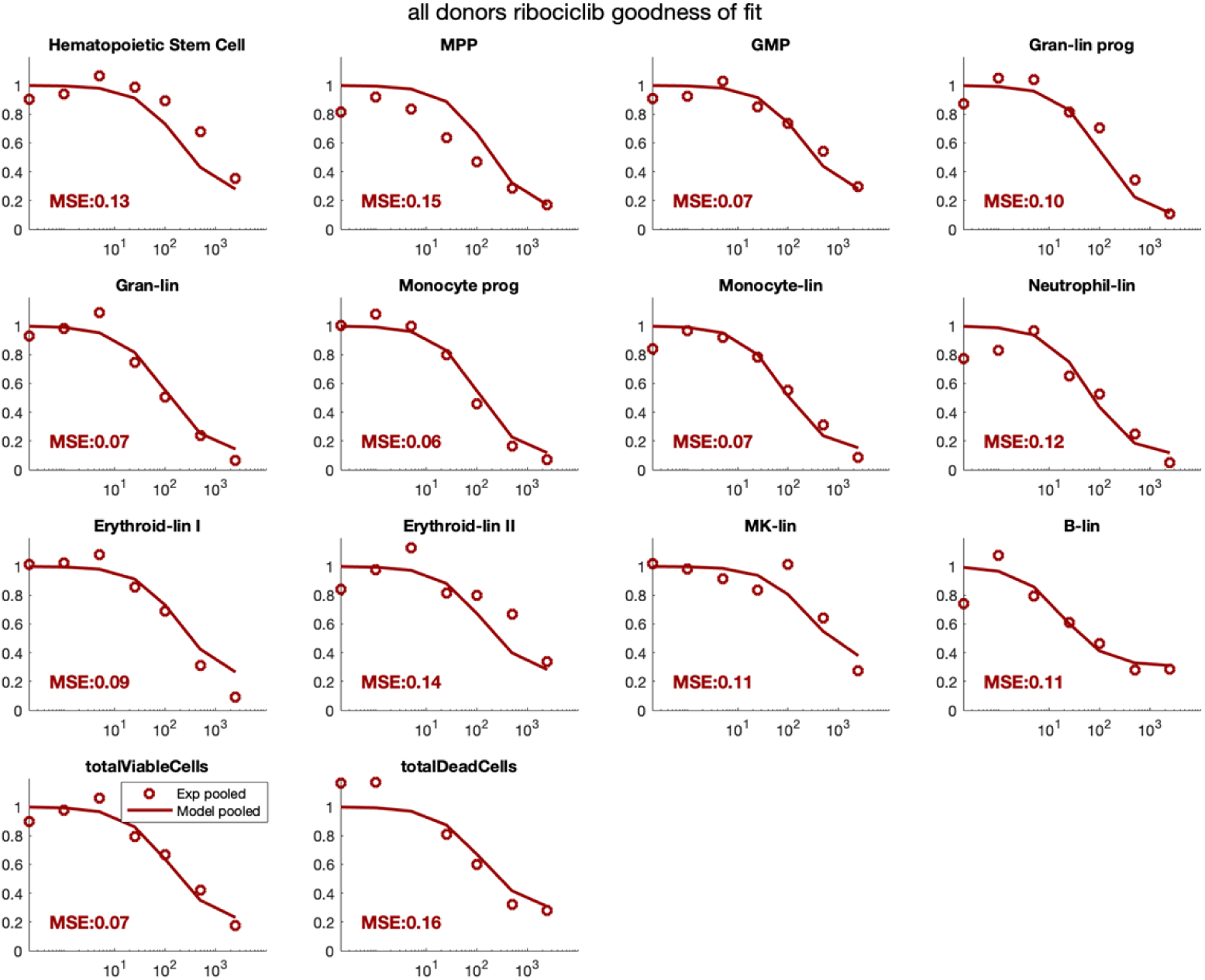
Goodness of fit across cell types for ribociclib. For each cell type and total live and viable cells, normalized cell counts from experimental data (open circles) and simulated results (solid line) are plotted against concentration (nM). Additionally, each plot includes the mean squared error of the difference between experimental and simulated data.

**Supplemental Figure 5.**
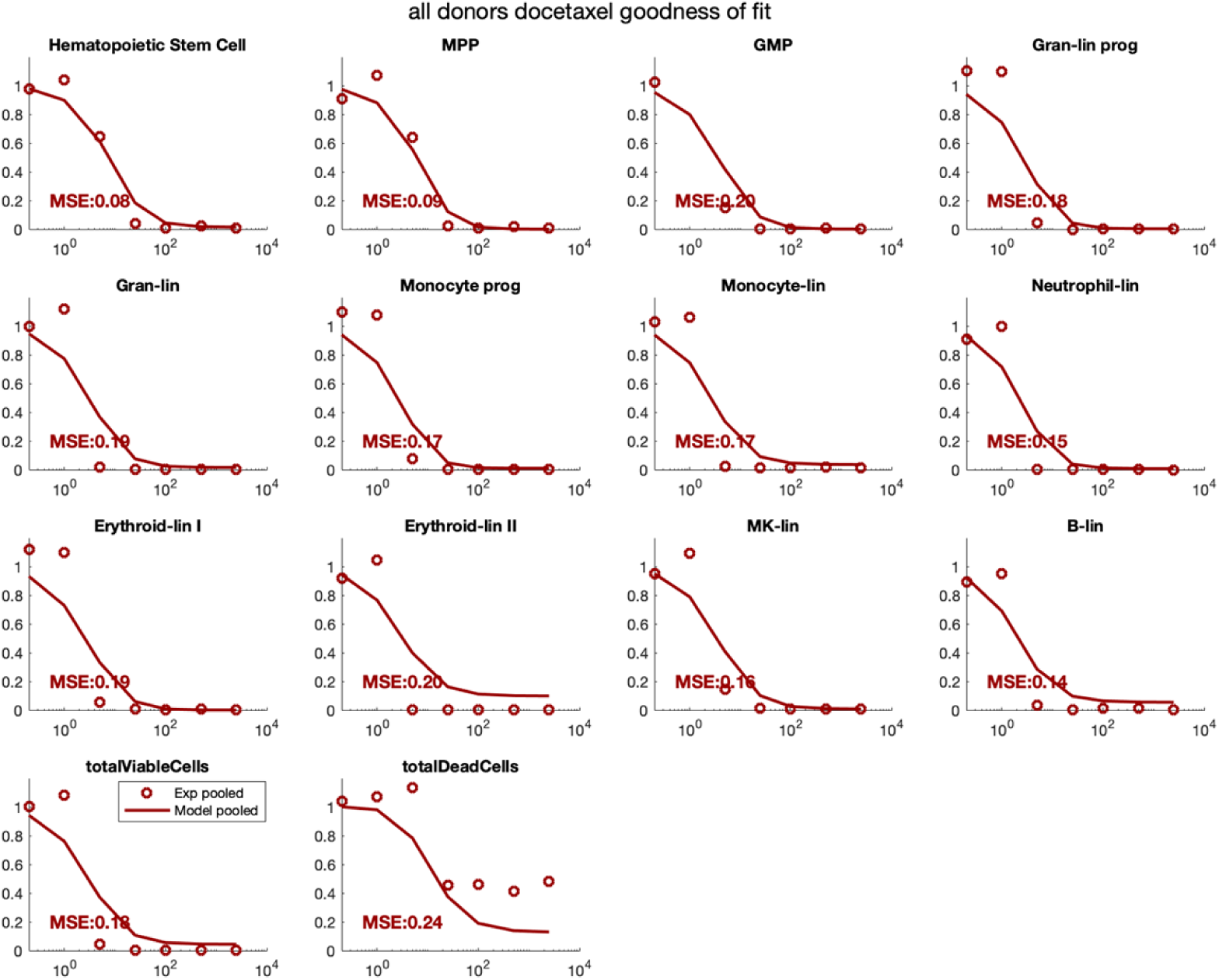
Goodness of fit across cell types for docetaxel. For each cell type and total live and viable cells, normalized cell counts from experimental data (open circles) and simulated results (solid line) are plotted against concentration (nM). Additionally, each plot includes the mean squared error of the difference between experimental and simulated data.

**Supplemental Figure 6.**
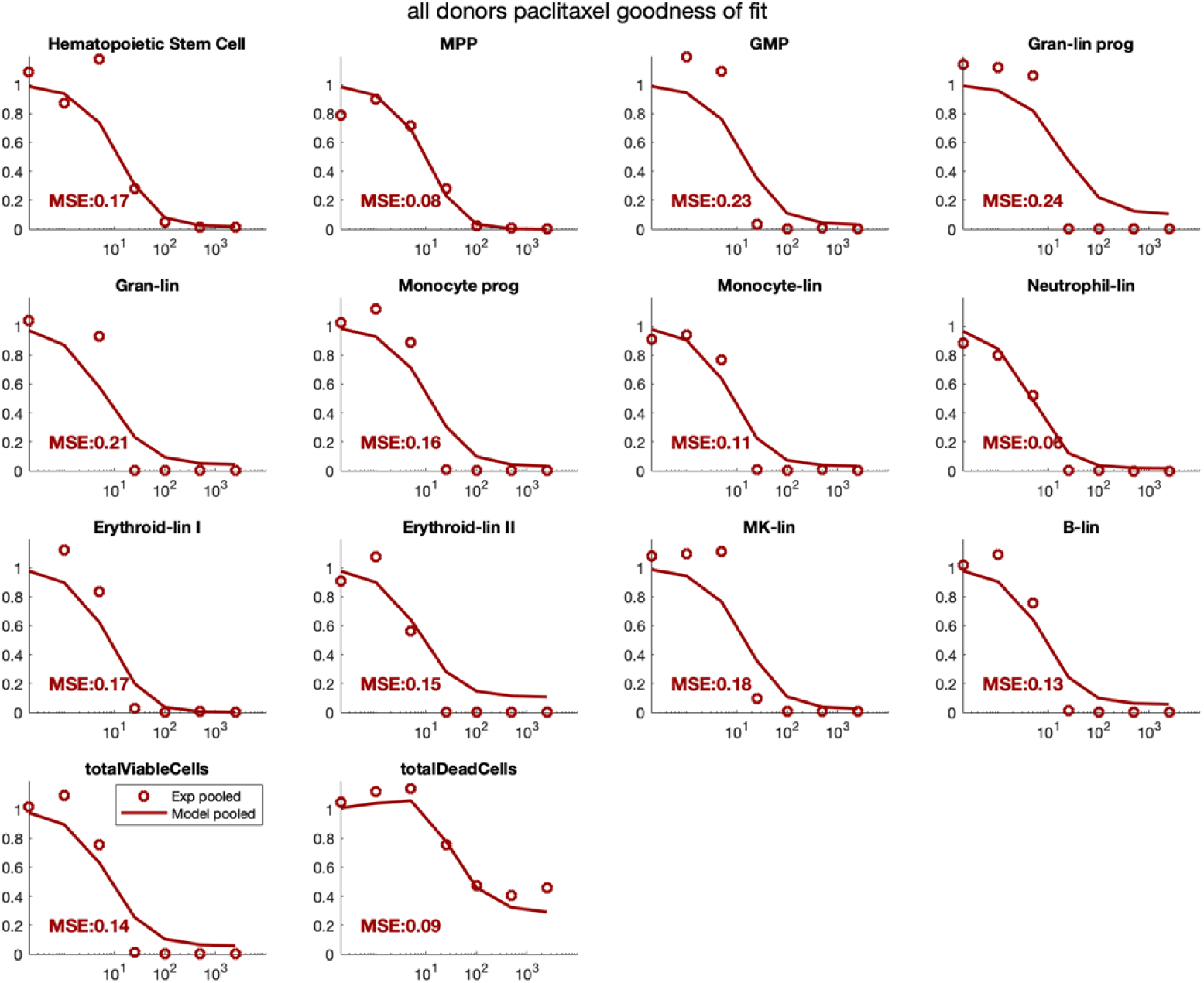
Goodness of fit across cell types for paclitaxel. For each cell type and total live and viable cells, normalized cell counts from experimental data (open circles) and simulated results (solid line) are plotted against concentration (nM). Additionally, each plot includes the mean squared error of the difference between experimental and simulated data.

**Supplemental Figure 7.**
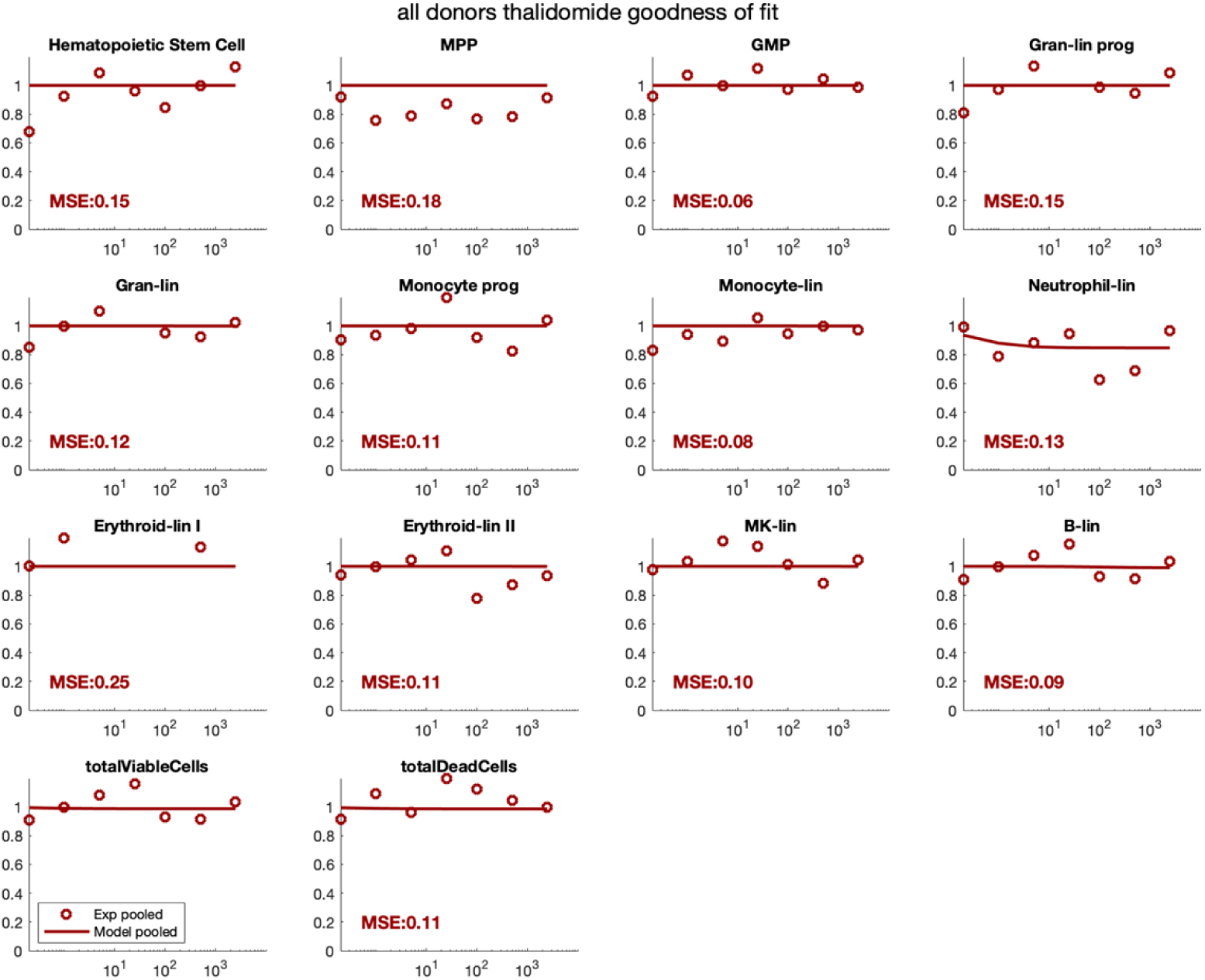
Goodness of fit across cell types for thalidomide. For each cell type and total live and viable cells, normalized cell counts from experimental data (open circles) and simulated results (solid line) are plotted against concentration (nM). Additionally, each plot includes the mean squared error of the difference between experimental and simulated data.

**Supplemental Figure 8.**
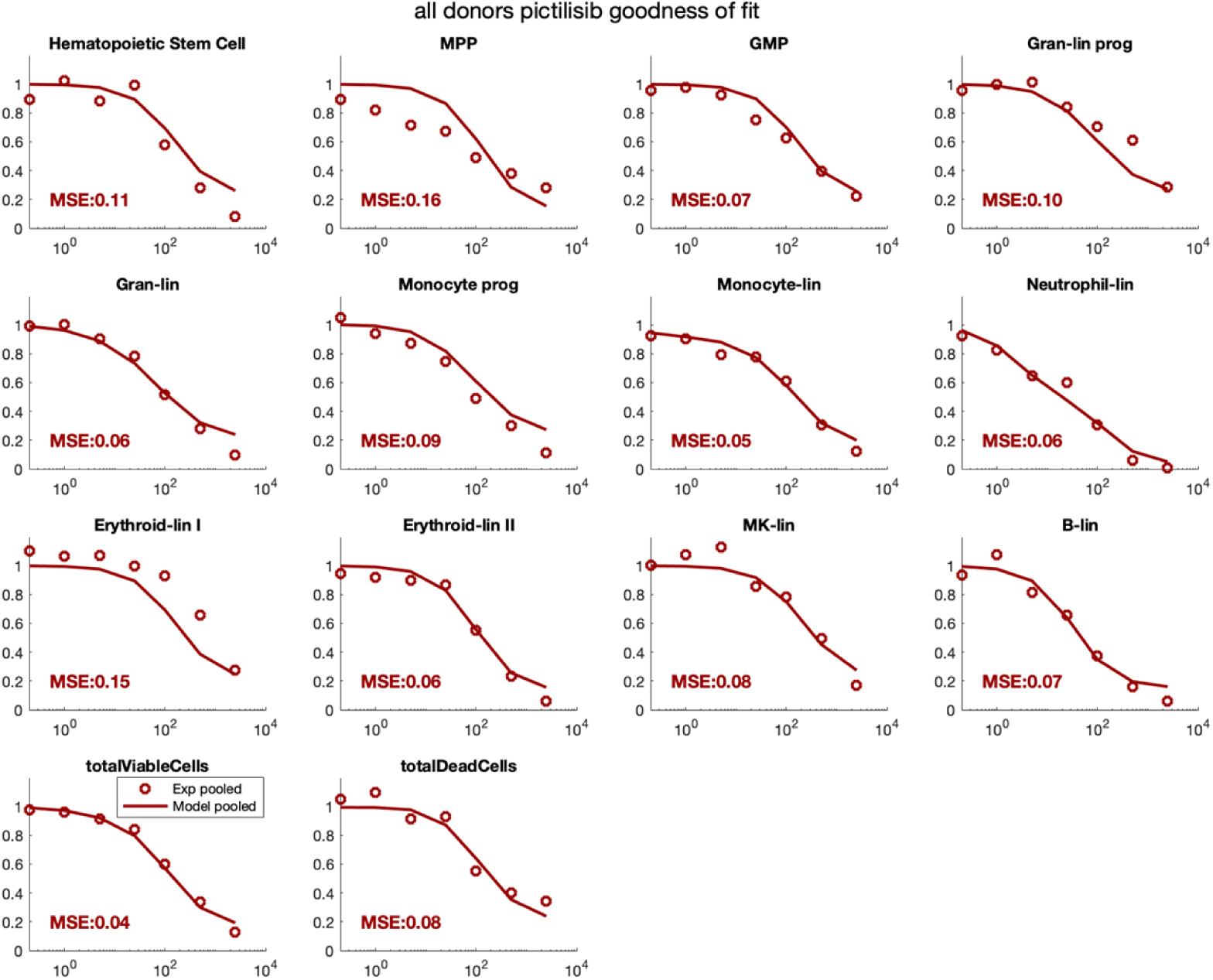
Goodness of fit across cell types for pictilisib. For each cell type and total live and viable cells, normalized cell counts from experimental data (open circles) and simulated results (solid line) are plotted against concentration (nM). Additionally, each plot includes the mean squared error of the difference between experimental and simulated data.

**Supplemental Figure 9.**
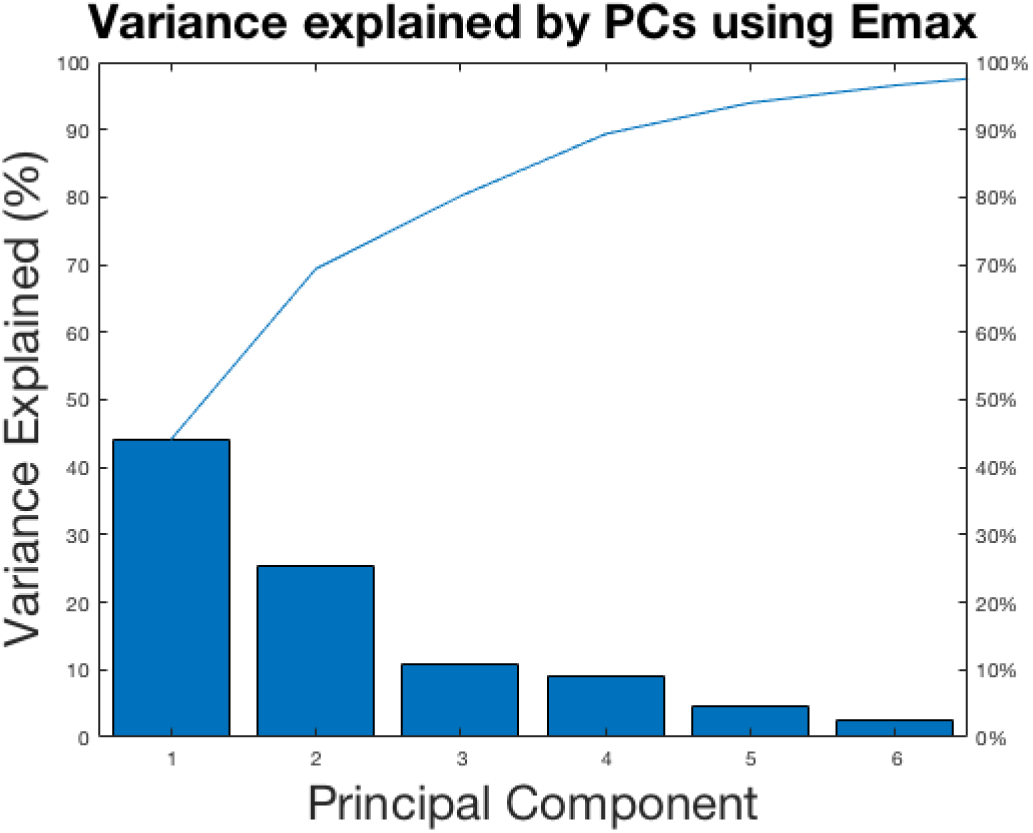
Percent variance explained by each principal component. The percent variance(left axis) and cumulative variance (right axis) are plotted against the top six components (x-axis).

**Supplemental Figure 10.**
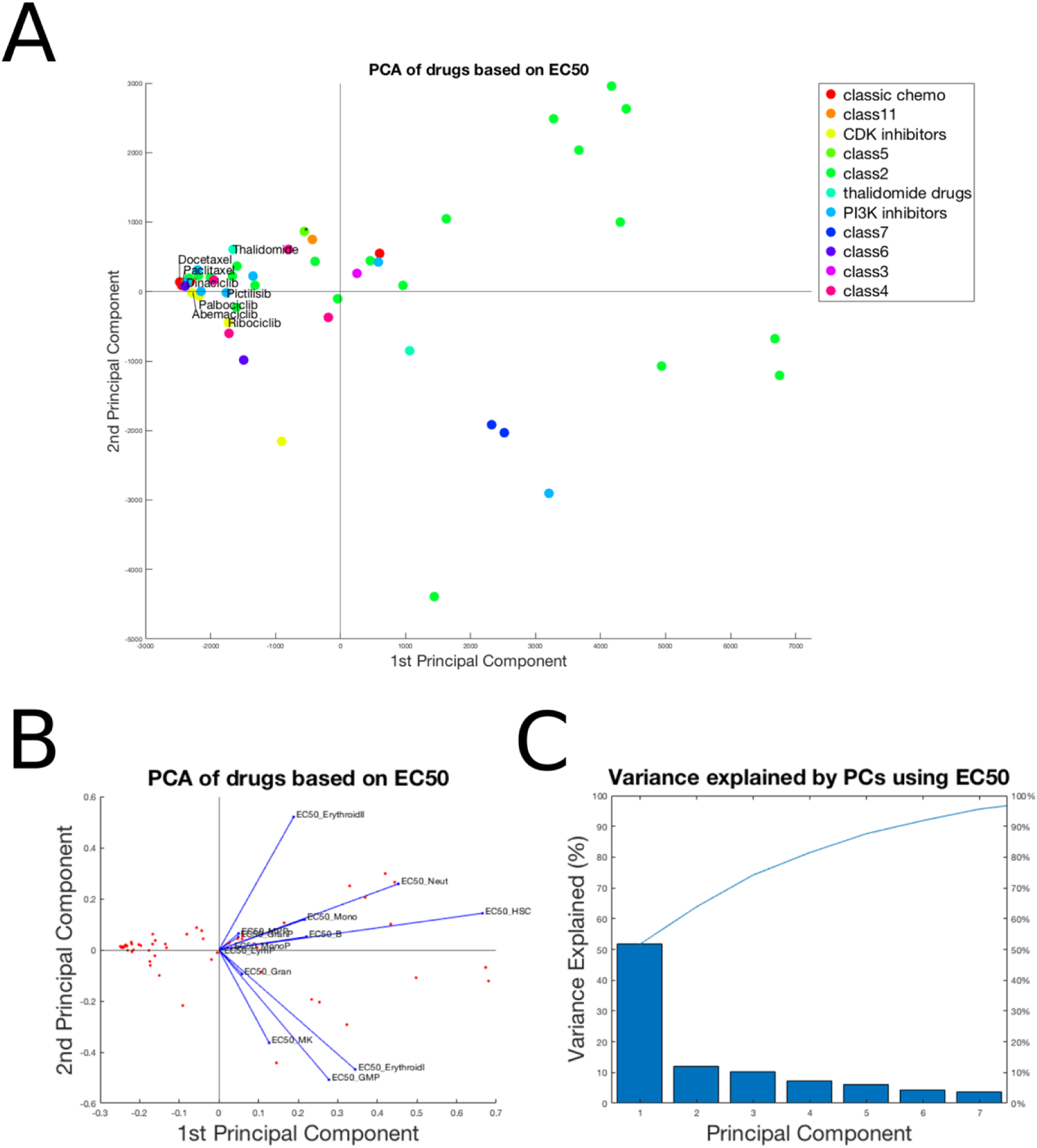
Principal component analysis of drugs based on the Emax of their effects. The 51 compounds are plotted in PCA space (**A**). Marker color corresponds to drug class. Drugs and variables contributing to the top two components are plotted in PCA space (**B**). Note: in figure **A** there is only one drug in class 5 and it is marked with an * to distinguish this compound from the remaining class 2 drugs. Variance explained by each principal component is plotted in (**C**).

**Supplemental Figure 11.**
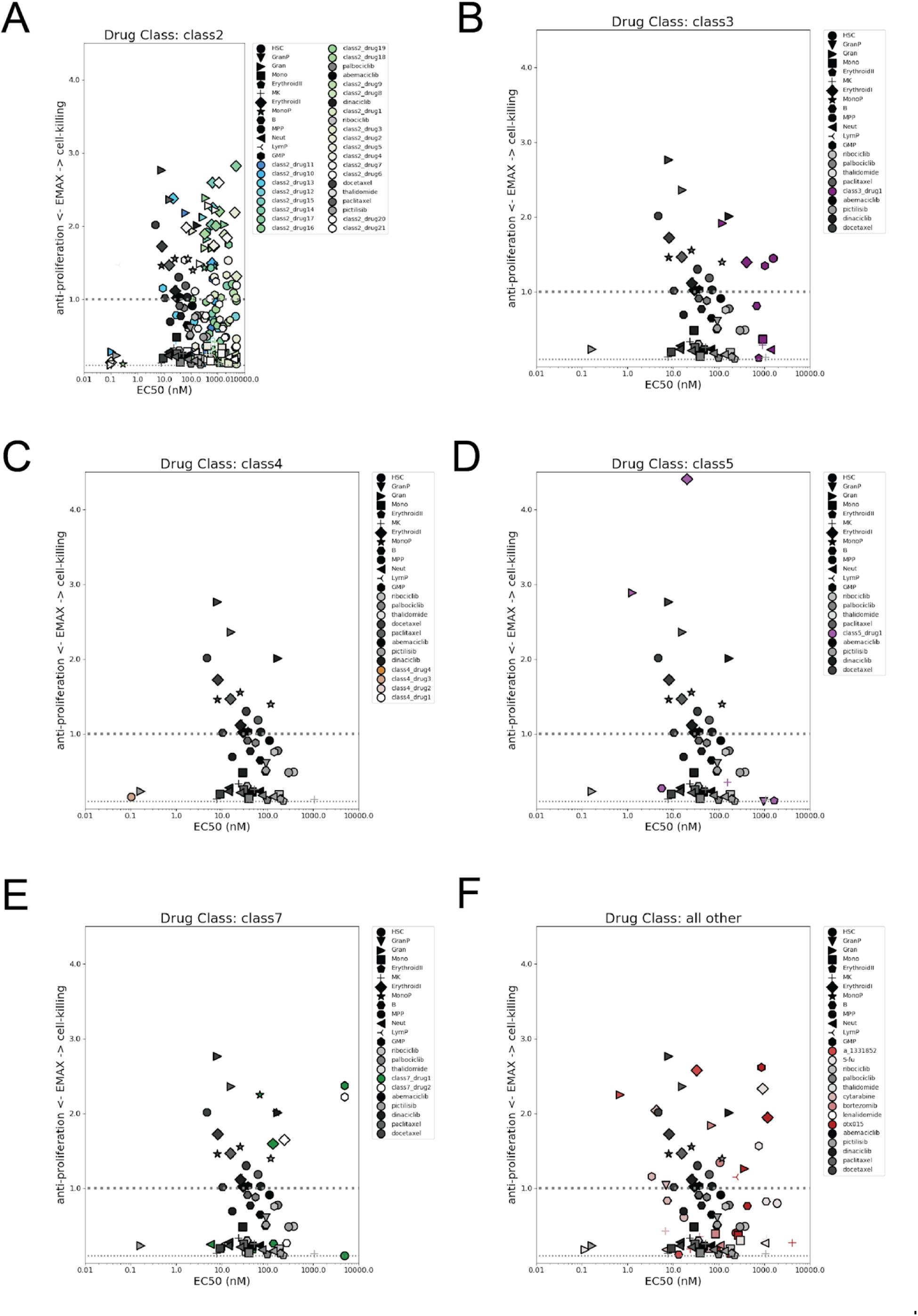
Mechanistic effects of developmental compounds per cell type compared to the sample set. Emax parameters per cell type are plotted against the EC50 values for each cell type. Marker shape represents cell type and marker shading represents either the sample set (black gradient, all figures) or drugs in class 2 (blues, **A**), class 3 (purples, **B**), class 4 (browns, **C**), class 5 (orchids, **D**), class 7 (green, **E**), or all other drugs (reds, **F**). The dashed line represents where Emax = 1.0.

## References

1. Barreto JN, McCullough KB, Ice LL, Smith JA. Antineoplastic Agents and the Associated Myelosuppressive Effects: A Review. Journal of Pharmacy Practice. 4 ed. SAGE PublicationsSage CA: Los Angeles, CA; 2014 Aug 20;27(5):440–6.

2. Portenoy RK, Itri LM. Cancer-related fatigue: guidelines for evaluation and management. Rehabilitation Oncology. 2001;19(2):32.

3. Craig M. Towards Quantitative Systems Pharmacology Models of Chemotherapy-Induced Neutropenia. CPT: pharmacomet syst pharmacol. John Wiley & Sons, Ltd; 2017 May 1;6(5):293–304.

4. Bodey GP, Buckley M, Sathe YS, Freireich EJ. Quantitative relationships between circulating leukocytes and infection in patients with acute leukemia. Am Coll Physicians. 1966 Feb;64(2):328–40.

5. Venkatakrishnan K, Friberg LE, Ouellet D, Mettetal JT, Stein A, Trocóniz IF, et al. Optimizing Oncology Therapeutics Through Quantitative Translational and Clinical Pharmacology: Challenges and Opportunities. Clinical Pharmacology & Therapeutics. John Wiley & Sons, Ltd; 2015 Jan 1;97(1):37–54.

6. Collins TA, Hattersley MM, Yates J, Clark E, Mondal M, Mettetal JT. Translational Modeling of Drug-Induced Myelosuppression and Effect of Pretreatment Myelosuppression for AZD5153, a Selective BRD4 Inhibitor. CPT: pharmacomet syst pharmacol. John Wiley & Sons, Ltd; 2017 Jun 1;6(6):357–64.

7. Pessina A, Albella B, Bayo M, Bueren J, Brantom P, Casati S, et al. Application of the CFU-GM assay to predict acute drug-induced neutropenia: an international blind trial to validate a prediction model for the maximum tolerated dose (MTD) of myelosuppressive xenobiotics. Toxicol Sci. 2003 Oct;75(2):355–67.

8. Friberg LE, Henningsson A, Maas H, Nguyen L, Karlsson MO. Model of Chemotherapy-Induced Myelosuppression With Parameter Consistency Across Drugs. Journal of Clinical Oncology. 2002;20(24):4713–21.

9. Quartino AL, Karlsson MO, Lindman H, Friberg LE. Characterization of Endogenous G-CSF and the Inverse Correlation to Chemotherapy-Induced Neutropenia in Patients with Breast Cancer Using Population Modeling. Pharm Res. Springer US; 2014 Jun 12;31(12):3390–403.

10. Sun W, Yu Y, Hoffman J, Turner NC, Cristofanilli M, Wang DD. Palbociclib exposure-response analyses in second-line treatment of hormone-receptor positive advanced breast cancer (ABC). Journal of Clinical Oncology. American Society of Clinical Oncology; 2017 May 30;35(15_suppl):1053–3.

11. Fornari C, O’Connor LO, Yates JWT, Cheung SYA, Jodrell DI, Mettetal JT, et al. Understanding Hematological Toxicities Using Mathematical Modeling. Clinical Pharmacology & Therapeutics. John Wiley & Sons, Ltd; 2018 Oct 1;104(4):644– 54.

12. Fornari C, O’Connor LO, Pin C, Smith A, Yates JWT, Cheung SYA, et al. Quantifying drug-induced bone marrow toxicity using a novel haematopoiesis systems pharmacology model. CPT: pharmacomet syst pharmacol. John Wiley & Sons, Ltd; 2019 Sep 11;:psp4.12459.

13. Li W, Lam MS, Birkeland A, Riffel A, Montana L, Sullivan ME, et al. Cell-based assays for profiling activity and safety properties of cancer drugs. Journal of Pharmacological and Toxicological Methods. Elsevier; 2006 Nov 1;54(3):313–9.

14. Morse DL, Gray H, Payne CM, Gillies RJ. Docetaxel induces cell death through mitotic catastrophe in human breast cancer cells. Mol Cancer Ther. American Association for Cancer Research; 2005 Oct 1;4(10):1495–504.

15. Weaver BA. How Taxol/paclitaxel kills cancer cells. Bement W, editor. MBoC. 2014 Sep 15;25(18):2677–81.

16. Hu J, Sun C, Bernatchez C, Xia X, Hwu P, Dotti G, et al. T-cell Homing Therapy for Reducing Regulatory T Cells and Preserving Effector T-cell Function in Large Solid Tumors. Clin Cancer Res. American Association for Cancer Research; 2018 Jun 15;24(12):2920–34.

17. Mauch P, Constine L, Greenberger J, Knospe W, Sullivan J, Liesveld JL, et al. Hematopoietic stem cell compartment: Acute and late effects of radiation therapy and chemotherapy. Int J Radiat Oncol Biol Phys. Elsevier; 1995 Mar 30;31(5):1319–39.

18. Marciniak-Czochra A, Stiehl T. Mathematical models of hematopoietic reconstitution after stem cell transplantation. In Model Based Parameter Estimation 2013 (pp. 191–206). Springer, Berlin, Heidelberg.

